# Microglia integration into human midbrain organoids leads to increased neuronal maturation and functionality

**DOI:** 10.1101/2022.01.21.477192

**Authors:** Sonia Sabate-Soler, Sarah Louise Nickels, Cláudia Saraiva, Emanuel Berger, Ugne Dubonyte, Kyriaki Barmpa, Yan Jun Lan, Tsukasa Kouno, Javier Jarazo, Graham Robertson, Jafar Sharif, Haruhiko Koseki, Christian Thome, Jay W. Shin, Sally A. Cowley, Jens C. Schwamborn

## Abstract

The human brain is a complex, three-dimensional structure. To better recapitulate brain complexity, recent efforts have focused on the development of human specific midbrain organoids. Human iPSC-derived midbrain organoids consist of differentiated and functional neurons, which contain active synapses, as well as astrocytes and oligodendrocytes. However, the absence of microglia, with their ability to remodel neuronal networks and phagocytose apoptotic cells and debris, represents a major disadvantage for the current midbrain organoid systems. Additionally, neuro-inflammation related disease modeling is not possible in the absence of microglia. So far, no studies about the effects of human iPSC-derived microglia on midbrain organoid neural cells have been published. Here we describe an approach to derive microglia from human iPSCs and integrate them into iPSC-derived midbrain organoids. Using single nuclear RNA Sequencing, we provide a detailed characterization of microglia in midbrain organoids as well as the influence of their presence on the other cells of the organoids. Furthermore, we describe the effects that microglia have on cell death and oxidative stress-related gene expression. Finally, we show that microglia in midbrain organoids affect synaptic remodeling and increase neuronal excitability. Altogether, we show a more suitable system to further investigate brain development, as well as neurodegenerative diseases and neuro- inflammation.

**Main Points:**

– Macrophage precursors can be efficiently co-cultured with midbrain organoids, they integrate into the tissue and differentiate into microglia in 3D.
– Organoids containing microglia have a smaller size and show a down-regulation of oxidative stress-related genes.
– Organoids co-cultured with microglia show differences in genes related to synaptic remodeling and action potential, as well as a more spontaneous action potential firing.

## Introduction

The human brain is a highly complex organ in terms of structure, molecular and cellular composition, making it a challenging target for research. Three-dimensional (3D) brain models have been recently developed to better mimic the spatial and functional complexity of the human brain. Over the last decade, different protocols were developed to generate either whole brain organoids (Lancaster *et al*., 2013; Lindborg *et al*., 2016) or discrete regions of the brain(Shi *et al*., 2012; Jo *et al*., 2016; Qian *et al*., 2016; Birey *et al*., 2017; Monzel *et al*., 2017). These models have proven to be suitable to model neurological disorders, including microcephaly (Lancaster *et al*., 2013), Batten disease (Gomez-Giro *et al*., 2019), Parkinson’s disease (PD) (Smits *et al*., 2019) and others (Choi *et al*., 2014; Qian *et al*., 2016). In fact, to better model PD recent efforts have focused on the development of PD patient specific midbrain organoids (Kim *et al*., 2019; Smits *et al*., 2019). Midbrain organoids contain spatially patterned groups of dopaminergic neurons, making them a suitable model to study PD. They consist of differentiated and functional neurons, which contain active synapses, as well as astrocytes and oligodendrocytes (Monzel *et al*., 2017; Smits *et al*., 2019). Moreover, they are able to recapitulate cardinal features of PD, including loss of dopaminergic neurons (Smits *et al*., 2019), and protein aggregation (Kim *et al*., 2019).

A major feature of neurological disorders is chronic inflammation (Bradburn, Murgatroyd and Ray, 2019; Tu *et al*., 2019). Previous studies have shown integration of iPSC-derived microglia in cerebral organoids (Muffat *et al*., 2016; Abud *et al*., 2017). Furthermore, innate differentiation of microglia within whole brain organoids has also been achieved(PR *et al*., 2018; Ramani *et al*., 2020). Those systems used iPSCs to generate their cerebral organoids, instead of a neuro- epithelial stem cell population, used normally to generate midbrain region-specific brain organoids. The current midbrain organoid system derived from neuro-epithelial stem cells– with ectodermal origin –lacks microglia, due to their mesodermal origin. The absence of microglia, with their ability to prune neuronal synapses as well as phagocytose apoptotic cells and debris represents a major disadvantage for the understanding of the physiological brain. Additionally, neuro-inflammation related disease modeling is not possible in a system that lacks microglia.

Microglia, the largest population of immune cells in the brain, are tissue-resident macrophages. In the adult brain they represent 5 to 15% of the adult brain cells, depending on the brain region (Thion, Ginhoux and Garel, 2018). Microglia have a unique ontogeny; they are derived from Yolk sac progenitors in a very early embryonic age (Ginhoux *et al*., 2010; Schulz *et al*., 2012; Li and Barres, 2017). Microglia have particular functions during brain development. Among others, they establish contacts with neural progenitors to support neurogenesis and proliferation (Choi *et al*., 2008; Ueno *et al*., 2013). In the adult brain, they interact with neurons, astrocytes and oligodendrocytes, and their major functions are maintenance of brain homeostasis and immune defense. They also interact with synapses, modulating neuronal activity, and perform synaptic pruning (Wake *et al*., 2009; Tremblay *et al*., 2011). They phagocytose apoptotic neurons, induce programmed cell death (Witting *et al*., 2000), guide sprouting blood vessels, and participate in neuronal maturation (Rymo *et al*., 2011). Chronic neuro-inflammation is one of the neuropathological characteristics of neurodegenerative disorders, including PD (Shabab *et al*., 2017).

Here, we describe the stable integration of functional human iPSC-derived microglia into midbrain organoids. This represents a significant advancement of the midbrain model, increasing its complexity and making a step forward from organoids to multi-lineage assembloids (Pasca, 2019; Marton and Pa ca, 2020). Moreover, microglia within assembloids express phagocytosis-related genes and release cytokines and chemokines, demonstrating relevant cellular communication abilities. Moreover, we demonstrate that assembloids display a reduction of cell death and oxidative stress-related genes in the system, compared to midbrain organoids without microglia. Furthermore, microglia seem to affect synapse remodeling, and lead to increased electrophysiology properties in neurons. Overall, we have established a stable and reproducible way to integrate functional microglia into midbrain organoids, leading to a next generation of midbrain organoid modelling. Our technology represents a step forward for the understanding and modulation of the complexity of the healthy and diseased human brain, with especial relevance for neuro-inflammatory conditions.

## Materials and Methods 2D cell culture

### iPSCs

Generation of iPSCs was performed as described in (Reinhardt *et al*., 2013). iPSCs (Table S1) where cultured in Matrigel^®^ (Corning, 354277) coated 6-well plates (Thermo Scientific, 140675), using Essential 8 Basal medium (Thermo Scientific, A1517001) supplemented with ROCK Inhibitor (Y-27632, Millipore, SCM075) for the first 24 hours after plating. The medium was exchanged on a daily basis. Confluence iPSCs (∼70-90%) slipt using Accutase^®^ (Sigma, A6964) and plated at around 300,000 cells per well. Neural progenitor cells needed to generated organoids were derived from iPSCs and maintained in culture as described previously (Smits *et al*., 2019; Nickels *et al*., 2020). iPSCs were also used to generate Macrophage precursors (van Wilgenburg *et al*., 2013) and further differentiate them into microglia as described previously (Haenseler, Sansom, *et al*., 2017).

### Microglia

50K macrophage precursors were plated per well in a glass bottom 96-glass bottom well plate (IBL Baustoff, 220.230.042). Cells were cultured with microglia differentiation medium (Advanced DMEM/F12 (Thermo Fisher, 12634010), 1x N2 (Thermo Fisher, 17502001), 1x Pen/Strep (Invitrogen, 15140122), 1x GlutaMax (Thermo Fisher, 35050061), 50 μ M 2-mercaptoethanol (Thermo Fisher, 31350-010), 100 ng/mL IL-34 (Peprotech, 200-34), 10 ng/mL GM-CSF (Peprotech, 300-03). Cells were kept in culture for 10 days, with a medium change every 3-4 days, and then fixed with 4 % formaldehyde (Millipore, 1.00496.5000) for immunostaining or used for a phagocytosis or MTT assay.

### 3D cell culture

#### Midbrain organoid generation and culture

Midbrain organoids generation is described in (Smits *et al*., 2019; Nickels *et al*., 2020). Shortly, 6,000 cells per well were seeded in an Ultra-Low Attachment 96-well Well plate (Merck, CLS3474) and kept under maintenance conditions (N2B27 medium supplemented with 0.2 mM Ascorbic acid (Sigma, A4544-100G), 3 μM M SB-431542 (Abcam, Smoothened Agonist (SAG, Stem cell technologies, 73412), , 2.5 μ M LDN-193189 (Sigma, SML0559)) for 2 days. After that, we started the pre- μ patterning (day 0 of dopaminergic differentiation) by removing SB and LDN from the medium. Two days after, CHIR concentration was reduced to 0.7 μM. On day 6 of dopaminergic μ differentiation, the medium was changed into maturation medium (N2B27 plus 0.2 mM Ascorbic acid, 10ng/mL Brain Derived Neurotrophic Factor, BDNF (Peprotech, 450-02), 10 ng/mL Glial-Derived Neurotrophic Factor, GDNF (Peprotech, 450-10), 1pg/mL TGF-β3 (Peprotech, 100-36E), 0.5 mM db cAMP (Sigma, D0627-5X1G), 10 μβ DAPT (R&D Systems, 2634/10) and 2.5ng/mL Activin A (Thermo Scientific, PHC9564)). Organoids were kept under static culture conditions with media changes every third day until day 15 of dopaminergic differentiation.

### Medium optimization for co-culture with macrophage precursors

Two batches of organoids and assembloids were used for the media optimization. Midbrain organoids from line K7 were used as controls. Midbrain organoids from line K7 were co-cultured with macrophage precursors, either from line K7, 163 or EPI. Therefore, graphs indicating “Midbrain organoids” refer to K7 midbrain organoids, and “Assembloids” refer to pooled results from K7 organoids co-cultured with K7, 163 and EPI microglia separately. For the media test, midbrain organoids and assembloids were cultured in the previously described maturation medium until day 15 of dopaminergic differentiation. Then, independently of the addition or not of macrophage precursors, the medium was kept the same, exchanged by microglia differentiation medium or co-culture medium (Table S2; Advanced DMEM/F12, 1x N2 supplement, 1x GlutaMAX^TM^, 50 μ M 2-mercaptoethanol, 100 U/mL Penicillin-Streptomycin, 100ng/mL IL-34, 10 ng/mL GM-CSF, 10 ng/mL BDNF, 10 ng/mL GDNF, 10 μM DAPT and 2.5 ng/mL Activin A). We kept the culture until day 35 of differentiation, fixed with 4 % formaldehyde (Millipore, 1.00496.5000) and proceeded to immunofluorescence staining.

### Co-culture of midbrain organoids with macrophage precursors

From day 15 of dopaminergic differentiation,– when the co-culture started - the culture medium of midbrain organoids and assembloids was replaced by co-culture medium containing 186,000 freshly harvested macrophage precursor cells per organoid. As described in the previous Methods and Results sections, K7 midbrain organoids were used as controls, and K7 organoids were separately co-cultured with macrophage precursors from lines K7, 163 or EPI. The “Assembloids” group on graphs represent the pooled results from organoids co-cultured with the K7, 163 and EPI microglia lines. After, the plate was centrifuged at 100 xg for 3 minutes to promote the attachment of the cells to the surface of the organoids. Medium was changed every 2-3 days and the system was kept for 20 or 70 days (until day 35 or 85 of dopaminergic differentiation, respectively). Then, organoids and assembloids were snap-frozen for sequencing and protein extraction or fixed with 4 % formaldehyde (Millipore, 1.00496.5000) for immunofluorescence staining.

### Phagocytosis assay

For immunofluorescence staining macrophage precursors were harvested and 30,000 cells per well were plated in 96-glass bottom well plates (IBL Baustoff, 220.230.042) and differentiated into microglia. At day 10, two Zymosan A (S. cerevisiae) BioParticles™ (Thermo Fisher, Z23373) per cell were added (60,000 particles / well). Then, cells were incubated for 30 minutes at 37C, washed with 1x PBS and fixed with 4 % formaldehyde (Millipore, 1.00496.5000). Then fixed samples were used for immunofluorescence staining. For live imaging (Video S1, S2 and S3), 100,000 cells per well were seeded in 8-well Nunc™ Lab-Tek™ Chamber Slides (Thermo Fisher, 177402PK) and immediately imaged.

### Viability assay (MTT)

50K macrophage precursors were plated per well in a 96-glass bottom well plate (IBL Baustoff, 220.230.042). After 10 days of microglia differentiation induction, 10ul of 5 mg/ml MTT (3- [4,5-dimethylthiazol-2-yl]-2,5-diphenyltetrazolium bromide (Sigma, M2128)) were added to each well. Cells were incubated for 4h at 37C. Then, the medium was aspirated and 100ul of DMSO were added to each well, pipetting vigorously in order to detach and disrupt the cells. Absorbance was measure at 570 nm. Results were compared to midbrain organoid medium (MOm).

### Immunofluorescence staining in 2D

Cells cultured in glass coverslips (150K cells/well) or 96-well imaging plates were fixed for 15 min with 4 % formaldehyde (Sigma, 100496 ) and washed 3x with PBS. Permeabilization was done by using 0.3% Triton X-100 in 1x PBS for 15 minutes. The cells were washed 3 times with PBS and blocked the cells with 3% BSA (Carl Roth, 80764) in PBS at room temperature. Cells were then incubated in a wet chamber, overnight (16h) at 4°C with the primary antibodies (diluted in 3% BSA + 0.3% Triton X-100 in 1xPBS). Cells were rinsed 3 times with PBS and further incubated with secondary antibodies diluted in 3% BSA + 0.3% Triton X-100 in 1x PBS for 1 hour at room temperature. After 3 more PBS washes, the plates were directly imaged and the coverslips were mounted in a glass slide with Fluoromount-G® (Southern Biotech, Cat. No. 0100-01). The antibodies used are listed in Table S3.

### Immunofluorescence staining in 3D

Midbrain organoids and assembloids were fixed with 4 % Formaldehyde overnight at 4 °C and washed 3 times with PBS for 15 min. Organoids were embedded in 4 % low-melting point agarose (Biozym, Cat. No. 840100) in PBS. 70 m sections sections were obtained using a vibratome (Leica VT1000 S). We selected sections coming from the center of the organoids and assembloids, rather than border sections. The sections were blocked with 0.5 % Triton X-100, 0.1 % sodium azide, 0.1 % sodium citrate, 2 % BSA and 5 % donkey serum in PBS for 90 min at room temperature. Primary antibodies were diluted in 0.1 % Triton X-100, 0.1 % sodium azide, 0.1 % sodium citrate, 2 % BSA and 5 % donkey serum and were incubated for 48 h at 4 °C in a shaker. After incubation with the primary antibodies (Table S3), sections were washed 3x with PBS and subsequently incubated with secondary antibodies (Table S3) in 0.05 % Tween-20 in PBS for 2 h at RT and washed with 0.05 % Tween-20 in PBS. Sections were mounted in Fluoromount-G mounting medium on a glass slide.

### Imaging

Qualitative images were acquired with a confocal laser-scanning microscope (Zeiss LSM 710). For quantitative image analysis, the Operetta CLS High-Content Analysis System (Perkin Elmer) was used to automatically acquire 25 planes per organoid section, with an spacing between planes of 1 µm. Images were modified with the ZEN blue Software. For live imaging (supplemental videos), the Cell Observer SD and the CSU-X1 Spinning Disc Unit (ZEISS) were used. 100 frames were acquired in each video using the ZEN blue software, during 3054.06s (163 line), 3053.27s (EPI line) and 3053.66s (K7 line). The videos were processed and modified with Adobe Premiere and Screenpresso softwares in order to obtain a representative time-line of the phagocytosis process. The shown videos represent 29.97 frames per second, resulting on a total number of 659.34 frames (163 line), 689.31 frames (EPI line) and 449.55 frames (K7 line).

### Image analysis

To measure the area of organoids and assembloids, we used ImageJ. We delimited the perimeter of the organoids and assembloids and used the “Measure” tool to obtain a pixel surface. Surfaces were then compared using Graphpad Prism and displayed as ‘fold change’ in graphs.

For immunofluorescence staining analysis, 3D images of midbrain organoids and assembloids were analyzed in Matlab (Version 2017b, Mathworks) following (Bolognin *et al*., 2019; Smits *et al*., 2019). The in-house developed image analysis algorithms facilitate the segmentation of Nuclei, neurons and microglia, obtaining as a result the positive pixel surface for a selected marker. To estimate the Iba1 positive cell number, we used the *regionprops* Matlab function in 8 sections coming from 4 assembloids from 2 different batches. This function detected different a total number of 310 IBA1^+^ events. We separated the Hoechst^+^ detection by ‘live nuclei” (bigger and less intense) and “pyknotic nuclei” (smaller and more intense due to the chromatin condensation). A total number of 776 live nuclei and 4178 pyknotic nuclei were detected. After that, we averaged the volume results for IBA1, live nuclei and pyknotic nuclei, and we added a new line in the main image analysis function to automatically divide the IBA1 Mask (pixel volume) by the averaged IBA1 volume number previously calculated. We did the same for the live nuclei, obtaining then an approximate IBA1^+^ cell number, and live nuclei number. We obtained the percentage of IBA1^+^ cells following this formula: (IBA1^+^ cell number /live nuclei number) x 100. The reason why we used the live nuclei number is that dead cells may lose the IBA1 expression.

### Western blot

Protein was extracted from midbrain organoids from 3 batches, and from assembloids with microglia from line K7, 163 and EPI from 3 batches. Protein extraction was done from 5 or 4 pooled organoids using the 1x RIPA buffer (Abcam, ab156034) containing 1x Phosphatase inhibitor cocktail (Merck Millipore, 524629-1ML) and 1x cOmplete™ Protease Inhibitor Cocktail (Sigma, 11697498001). The lysates were sonicated in the Bioruptor (Diagenode) for 10 cycles of 30 sec on and 30 sec off. The amount of protein was measured using the Pierce BCA Protein Assay Kit (Thermofisher, 23227) and the BIOTEK Cytation 5 Imaging reader. 30 μg of protein was used from each sample in the western blot. 6x loading buffer containing 0.375M Tris (MW 121.14 g/mol), 9% SDS, 50% Glycerol, 0.2% Bromphenolblue, 0.3M DTT was added in each sample and boiled at 95oC for 5 min. The protein samples were loaded into precast polyacrylamide gels (Thermofisher, NW04120BOX). The iBlot2 device from Invitrogen was used for transferring the proteins from the gel to PVDF membranes (Thermofisher, IB24001).

The membranes were blocked in 1xPBS containing 0.2% Tween-20 and 5% Milk in for 60 min at RT. Primary antibodies were incubated in 1XPBS containing 0.02% Tween-20, 5% BSA at 4°C overnight. Secondary antibodies were incubated for 60 min at RT in the same buffer as the one for primary antibodies. Enhanced fluorescent signal was detected in the LI-COR OdysseyFc imaging system. Band quantifications were performed with ImageJ and statistics were run using GraphPad Prism. A Mann-Whitney test was run to compare the midbrain organoid against the assembloid group.

### RT-PCR

Between one and three million microglia cells were used per RNA extraction. We used the RNeasy Mini Kit (Qiagen) as well as DNase I Amplification Grade (Sigma-Aldrich) to isolate RNA. After conducting reverse395 transcription by following the protocol of the High Capacity RNA to DNA Kit (Thermo Fisher Scientific), RT–PCRs were performed using GreenTaq polymerase and 50ng of cDNA per reaction. An initial denaturing step, 5 min at 95 °C, 40 cycles of denaturation for 30 s at 95 °C, annealing for 45 s at 55 °C (for Iba-1, CD68, and TMEM119) or 61 °C (for RPL37A and P2RY12), extension for 30 s at 72 °C and a final extension for 5 min at 72 °C. The used primers are listed in Table S4.

### Cytokine and chemokine release assay

Cytokine and chemokine measurements were performed using the Human XL Cytokine Discovery Luminex® Performance Assay (RD Systems, #FCTSM18). We collected supernatants from three midbrain organoid and assembloid batches and three biological replicates (microglia lines K7, 163 and EPI). When values were too low to be detected, they appeared as ‘out of range’. In order to consider them statistically, they were assigned the lowest measured value of the standard curve for that metabolite. The statistics were run with three batches and three cell lines for the assembloid group (midbrain organoids vs assembloids K7, 163 and EPI).

### Metabolomics

For the extracellular metabolomics analysis, we used snap frozen media from 20 days of co-culture old midbrain organoids or assembloids after 48h of culture. We also incubated co-culture media, not in contact with organoids, as a control. 3 organoids or assembloids were used per batch, 3 batches were analyzed, and the 3 different cell lines were used for the assembloid group. From the measured results, the control (basal medium) was subtracted in order to discriminate secreted (positive numbers) against uptaken (negative numbers) metabolites.

### Polar metabolite extraction, derivatization, and GC-MS measurement

Extracellular metabolites from media samples were extracted using a methanolic extraction fluid (5:1, methanol/water mixture, v/v). The water fraction contained two internal standards Pentanedioic acid-D6 (c = 10 μg/mL; C/D/N Isotopes Inc.) and [UL-13C5]-Ribitol (c = 20 μL of medium was added to 240 μL ice-cold extraction l ice-cold chloroform, the mixture was shaken for 5 min at 4 °C. For fluid. After adding 100 phase separation, 100 μ l water were added and vortexed for 1 min. Then, the mixture was centrifuged at 21,000 xg for 5 min at 4 °C. 250 μ of the polar (upper) phase was transferred to GC glass vial with micro insert (5-250 μL) and evaporated to dry under μ vacuum at -4 °C.

Metabolite derivatization was performed by using a multi-purpose sample preparation robot (Gerstel). Dried medium extracts were dissolved in 30 μ \l pyridine, containing 20 mg/mL methoxyamine hydrochloride (Sigma-Aldrich), for 120 min at 45 °C under shaking. After adding 30 l N-methyl-N-trimethylsilyl-trifluoroacetamide (Macherey-Nagel), samples were incubated μ for 30 min at 45 °C under continuous shaking.

GC-MS analysis was performed by using an Agilent 7890A GC coupled to an Agilent 5975C inert XL Mass Selective Detector (Agilent Technologies). A sample volume of 1 μ into a Split/Splitless inlet, operating in split mode (10:1) at 270 °C. The gas chromatograph was equipped with a 30 m (I.D. 0.25 mm, film 0.25 μm) DB-5ms capillary column (Agilent J&W GC Column) with 5 m guard column in front of the analytical column. Helium was used as carrier gas with a constant flow rate of 1.2 ml/min. The GC oven temperature was held at 90 °C for 1 min and increased to 220 °C at 10 °C/min. Then, the temperature was increased to 280 °C at 20 °C/min followed by 5 min post run time at 325 °C. The total run time was 22 min. The transfer line temperature was set to 280 °C. The MSD was operating under electron ionization at 70 eV. The MS source was held at 230 °C and the quadrupole at 150 °C. Mass spectra were acquired in full scan mode (m/z 70 to 700).

### Data normalization and data processing

All GC-MS chromatograms were processed using MetaboliteDetector, v3.220190704 (REF). Compounds were annotated by retention time and mass spectrum using an in-house mass spectral library. The internal standards were added at the same concentration to every medium sample to correct for uncontrolled sample losses and analyte degradation during metabolite extraction. The data set was normalized by using the response ratio of the integrated peak area_analyte and the integrated peak area_internal standard. The results correspond to triplicates from three co-cultured batches and three biological replicates (midbrain organoids against assembloids with microglia from lines K7, 163 and EPI).

### Patch clamp

Passive and active electrophysiological properties of cells in assembloids (with microglia from line 163) and midbrain organoids were characterized by whole-cell patch-clamp recordings in voltage and current clamp.

Each organoid was transferred from the incubator to a submerged type recording chamber with constant perfusion of carbogen-buffered artificial cerebrospinal fluid (ACSF) at 32°C. The ACSF contained (in mM): 124 NaCl, 3 KCl, 1.8 MgSO4, 1.6 CaCl2, 10 glucose, 1.25 NaH2PO4, 26 NaH2CO3 with an osmolarity of 295 mOsm/l. The organoid was fixated between a large diameter pipette and a custom made platinum harp. Cells were visualized using phase-contrast on an upright BX51 microscope (Olympus, Hamburg, Germany) with a 60x water-immersion objective. Recording electrodes were pulled using borosilicate glass on a Flaming/Brown P-97 Puller (Sutter Instrument, Novato, CA, USA) to yield a resistance of 3–6 ΩM . The electrode solution contained (in mM): 126 potassium gluconate, 4 KCl, 10 HEPES, 0.3 EGTA, 4 MgATP, 0.3 Na2GTP, and 10 phosphocreatine adjusted to pH 7.2 using KOH and to 288 mOsm/l by adding sucrose. Recordings were obtained in voltage and current-clamp mode with an ELC- 03XS amplifier (NPI electronic, Tamm, Germany). Signals were low-pass filtered at 3 kHz and digitized with 20 kHz using a Micros 1401MKII AC-converter (CED, Cambridge, UK). Data were collected using the Signal 4.10 software (CED). Voltages were not corrected for the calculated liquid-junction potential of +14.5 mV. Test pulses of -50 pA and 100 ms were applied regularly to control for changes in series resistance.

Putative neurons were visually identified by their size and shape. After obtaining the whole-cell configuration, the resting membrane potential was determined in current clamp and the cell subsequently stabilized at -70 mV by continuous current injection. To assess active and passive membrane properties, hyper- and depolarizing current steps were injected at increments of 10 pA and of 500 ms length starting from -50 pA. Passive parameters were assessed by the smallest negative current step that yielded a constant plateau potential, which varied due to the heterogeneity of input resistances. To measure maximum firing rates, cells were depolarized by voltage steps of increasing amplitudes until APs started to fail due to sodium channel inactivation (up to +500 pA depending on input resistance). Action potential waveforms characteristics were analyzed for the first action potential that fired 50 ms after the onset of current injections. The voltage threshold was defined as the potential at which the rising slope exceeded 5 mV/ms and amplitudes were measured from threshold to peak. In voltage-clamp mode, cells were held at -70 mV and positive voltage steps of 300 ms duration and 10 mV increments were applied every 5 s. Maximal voltage was +30 mV. The power of voltage-gated cation channels (predominantly sodium) was quantified at -30mV and used for statistical comparison. Data analysis was performed using Stimfit (Guzman, 2014) and custom-written Phyton routines. Values were tested for Gaussian distribution by D’Agostino-Pearson omnibus normality test. Unpaired t-tests were used to assess statistical significance in normally distributed data and Mann-Whitney tests for non-normally distributed data (indicated as p in Figures). Outliers deviating 2.5 standard deviations were excluded from statistical analysis but indicated in the Figures.

The presence of microglia in the recorded assembloids was confirmed using immunohistochemical staining and confocal imaging. Organoids were fixed in 4% formaldehyde and stained for nuclei (Fluoroshield with DAPI, Sigma-Aldrich), neuronal markers markers (MAP2, Sigma-Aldrich Chemie GmbH), and microglia (anti-Iba1, WAKO Chemicals). Imaging was carried out on an A1 Nikon confocal microscope at the Nikon Imaging Center Heidelberg, Germany.

### Statistical analyses

First, Gaussian distribution was evaluated by performing D’Agostino & Pearson omnibus normality test. According to this distribution, either a 1way ANOVA or a Kruskal-Wallis test with a Dunnett’s test for multiple comparisons were used to evaluate statistical significance. For the pooled results (organoids against assembloids), gaussian distribution was also tested. Depending on the outcome, an unpaired t-test or Mann-Whitney test was used to assess the difference between groups. Outliers deviating 2.5 standard deviations were excluded from statistical analysis but indicated in the Figures. Cells that showed a resting membrane potential above -40 mV and action potentials shorter than 50 mV and wider than 3 ms in half-width were excluded from analysis, assuming that these cells were either not fully matured neurons or recording conditions were poor. The presence of microglia in the recorded organoids was confirmed using immunohistochemical staining and confocal imaging. For the image analysis, a 2way ANOVA, Tukey’s multiple comparisons test was performed to evaluate statistical significance. Data is presented as mean ± SEM. All analyses were performed with three different biological replicates (assembloids microglia from three different cell lines).

### Multi-electrode array (MEA)

48-well MEA plates (Axion, M768-tMEA-48B-5) were coated as follows: 24 h incubation with 0.1 mg/ml poly-D-lysine (Sigma, P7886) followed by a 24 h incubation with 1 mg/ml laminin (Sigma, L2020). At day 20 of co-culture (or 35 days of dopaminergic differentiation), midbrain organoids and assembloids were transferred to 48-well pre-coated MEA plates. Midbrain organoids and assembloids were placed in the center of the wells, and left in the incubator for 25 minutes to ensure the maximum media evaporation and adherence of the organoids and assembloids to the well bottom. Then, we added 25 ul of Geltrex (Invitrogen, A1413302) to the top, to avoid the detachment from the wells. Data was acquired with an Axion Maestro Multiwell 768-channel MEA System (Axion Biosystems) and using the Axis software (Axon Biosystems).

Data was exported and analyzed using the available R script following a published method ((Modamio *et al*., 2021) see ‘Resource availability’).

### Single-nuclei RNA sequencing (snRNAseq)

Five snap-frozen organoids or assembloids from one batch per condition were used to perform snRNAseq.

### Data generation

Whole frozen organoids and assembloids (with microglia from lines K7, 163 and EPI) were dissociated for generating single-nuclei gene-expression libraries. The following steps were performed on ice using chilled and freshly prepared buffers. In brief, organoids and assembloids were gently dissociated using 500 mL of 0.1X Lysis Buffer (10 mM Tris-HCl pH 7.4, 10 mM NaCl, 3 mM MgCl2, 0.1% Tween-20, 0.1% Nonidet P40 Substitute, 0.01% digitonin, 1% BSA, nuclease-free water) by pipetting 10X with a wide-pore 1000 μL pipette, followed by a 10 min incubation period on ice. This was repeated for 2 cycles. To reduce batch effects and increase the number of nuclei per experiment, material from 4 different organoids or assembloids were a new 40-μM strainer. The same procedure was done with 20- μL at a time, using M (Sysmex, AN777717) and 10-μM (Sysmex, AP275603) strainers, followed by centrifugation for 5 min at 500 xg, at 4°C.

Nuclei were then stained using Hoechst 33342 (1:2000) for 5 min at room temperature and counted on the Countess II.

Using Diluted Nuclei Buffer (10x Genomics, 2000153) each sample was adjusted to a L. We generated one library for each sample, aiming for 6000 nuclei. Single-nuclei experiments were performed using the 10x Genomics Next GEM Single Cell 5′ Library Kit v1.1 (1000168) to encapsulate nuclei and amplify cDNA, to generate sequencing libraries. Each library was barcoded using i7 barcodes provided by 10x Genomics. cDNA and sequencing library quality and quantity were determined using Agilent’s High Sensitivity DNA Assay (5067-4626) and KAPA (KK4824). Final libraries were pooled, loaded on two lanes of the Illumina’s HiSeq X and sequenced in 150PE mode.

### Count matrix generation

Following steps are performed on default settings if not otherwise specified. Single-nucleus libraries were demultiplexed based on their i7 index sequences and for each library mapping to the human genome was performed using the Cell Ranger 4.0.0 software and human reference GRCh38 3.0.0, both provided by 10x Genomics. Next, count matrix files for each sample were generated using cellranger count.

### Data pre-processing

Since the material coming from assembloids with the EPI microglia line was lost, count matrixes for the assembloids with K7 and 163 microglia, as well as the midbrain organoid control were uploaded and Seurat Objects were created. Data was preprocessed as decribed by https://satijalab.org/seurat/v3.2/pbmc3k_tutorial.html. An independent data quality control was performed of all three objects, by checking levels of ribosomal genes, and by removing cells with a high mitochondrial gene fraction (Lake *et al*., 2019; D and JJ, 2021). Moreover, cells containing less than 100 or more than 5000 (100 < RNA n Feature < 5000) genes were removed from the analysis, being considered empty droplets or doublets, respectively. After quality assessment all three objects were combined, log normalized, scaled, and a linear dimensional reduction was performed (PCA). Dimensionality was assessed (20) and cells were clustered with a resolution of 0.5. A non-linear dimensional reduction using “umap” was performed, and cluster markers were assessed. Cluster names were defined by their characteristic marker gene expression.

### Cell type identification

Cell clustering was performed based on the top 20 principal components using Louvain algorithm modularity optimization with a resolution of 0.5. UMAP was used for cell cluster visualization (Becht *et al*., 2018). The distinct cell clusters were identified in the UMAP plot. For cell type identification, binarized gene list across cell types from La Manno et al., 2016 was used. This list of genes comprises information about the marker genes in a binarized manner, where 1 means that gene is marking a specific cell population and 0 means that it cannot be considered as a marker gene. For more details on how this list is generated please refer to La Manno et al., 2016. Expression of each cluster defining gene was overlapped with the marker gene (1) in the La Manno marker matrix. Cell types were assigned based on the highest number of major cluster marker genes being expressed in the respective clusters. Cellular subtypes were verified and grouped in neuronal identity clusters based on neuronal marker gene lists (Table S5). (Chen *et al*., 2016; Smits *et al*., 2020). Moreover manual marker verifications were done using known marker genes for each cluster. For instance, RGLs expressed *SLC1A3 (Glast*). MidNESCs expressed midbrain markers *SHH*, *LMX1A* and *FOXA2* and stem cell markers *SOX2* and *MSI1* (*Musashi*). PROG highly expressed *VIM (Vimentin*) and lower levels of stem cell markers. NB were positive for young neuronal markers *DCX* (*doublecortin*) and *STMN1* (*sthaminin*) without expressing mature neuronal markers. YDN&CN still expressed *DCX* and *STMN1* but also some mature neuronal markers for synapses such as *SYP (synaptophysin*) and subtype specification markers such as *TH* (dopaminergic neurons) and *SLC18A3* (cholinergic neurons). Mature neurons expressed low levels of *STMN1* and *DCX* but high levels of *SYP*. Subtype specification revealed that mDN(A10)&GaN&GlN expressed high levels of genes that define neurons from the ventral tegmental area (VTA). Besides, cells expressing dopaminergic marker *TH* also expressed the marker for A10 DN *CALB1*, as well as *ADCYAP1*. Moreover, the glutamatergic *SLC18A1* and GABAergic *GAD1* transporters were highly expressed in these neurons. Last mDN(A9)&SN were qualified by high amounts of *TH* and KCNJ6 (*Girk2)* as well as DN defining synaptic markers *ROBO1*. Moreover, also serotonergic transporters *SLC*18A2 were expressed in that cluster. Microglial cells expressed *IBA1.* Cell type proportions were calculated by counting the cells of each cluster. A spearman correlation and a heatmap clustering (pheatmap) were performed on the average expression of clusters defining genes. Because of a substantial sample loss during sample processing, that affected the microglia cell count,. 26 microglial cells were detected (Table S6). The microglial cluster, 26 cells, was subsetted and an independent analysis was performed. The top 100 genes defining the microglial cluster are shown in a heatmap. Moreover, the average gene expression across all microglial cells was exported, and representative genes (Galatro et al. 2017). are presented in a violin plot. Significances were calculated based on a one sample Wilcoxon test *p<0.05. The same was done for genes involved in the phagocytic pathway. Cytokines and chemokines were represented in a heatmap.

### Differential expressed gene and enrichment analysis

The most variable features were identified and a heatmap clustering (pheatmap) was performed on the average expression of the top 100 most variable genes. DEG were identified between assembloids and midbrain organoids. This was done for all the cell types together and each cell type independently. Data of the DEG lists is represented in heatmaps (top 100), or underwent metacore enrichment analysis. Metacore analysis was based on the following arbitrary threshold p<0.05, adj p<0.5. Moreover, Venn diagrams were formed using nVenn http://degradome.uniovi.es/cgi-bin/nVenn/nvenn.cgi. Genes involved in the enriched pathways are shown separately in box plots. Statistics were performed using a Wilcoxon test p<0.05*.

## Results

### hiPSC-derived microglia express specific markers and are functional

The work presented here is based on the use of quality controlled hiPSC lines (lines: K7, 163, EPI, table S1) from three different healthy individuals that express the pluripotency markers SSEA-4, OCT-4, TRA-1-60, NANOG, TRA-1-81 and SOX2 (Figure S1A). We started by deriving macrophage precursors from hiPSCs that were then differentiated into microglia for 10 days, as previously described (Haenseler, Zambon, *et al*., 2017). hiPSC-derived microglia in monoculture expressed macrophage-specific markers, including IBA1, PU1 and CD45, and microglia markers, notably TMEM119 and P2RY12, as detected by immunostaining (Fig. 1A). Moreover, PCR confirmed that microglia also expressed the macrophage specific genes *IBA1* and *CD68*, and the microglia-specific genes *TMEM119* and *P2RY12* (Fig. 1B). After confirming the microglia cell identity, we next assessed their phagocytic capacity. Incubation of microglia with Zymosan particles showed the cells’ ability in phagocytosing these particles, detectable within their cell bodies (Figure 1C, Figure S1B, Video S1, S2 and S3).

**Figure 1.**
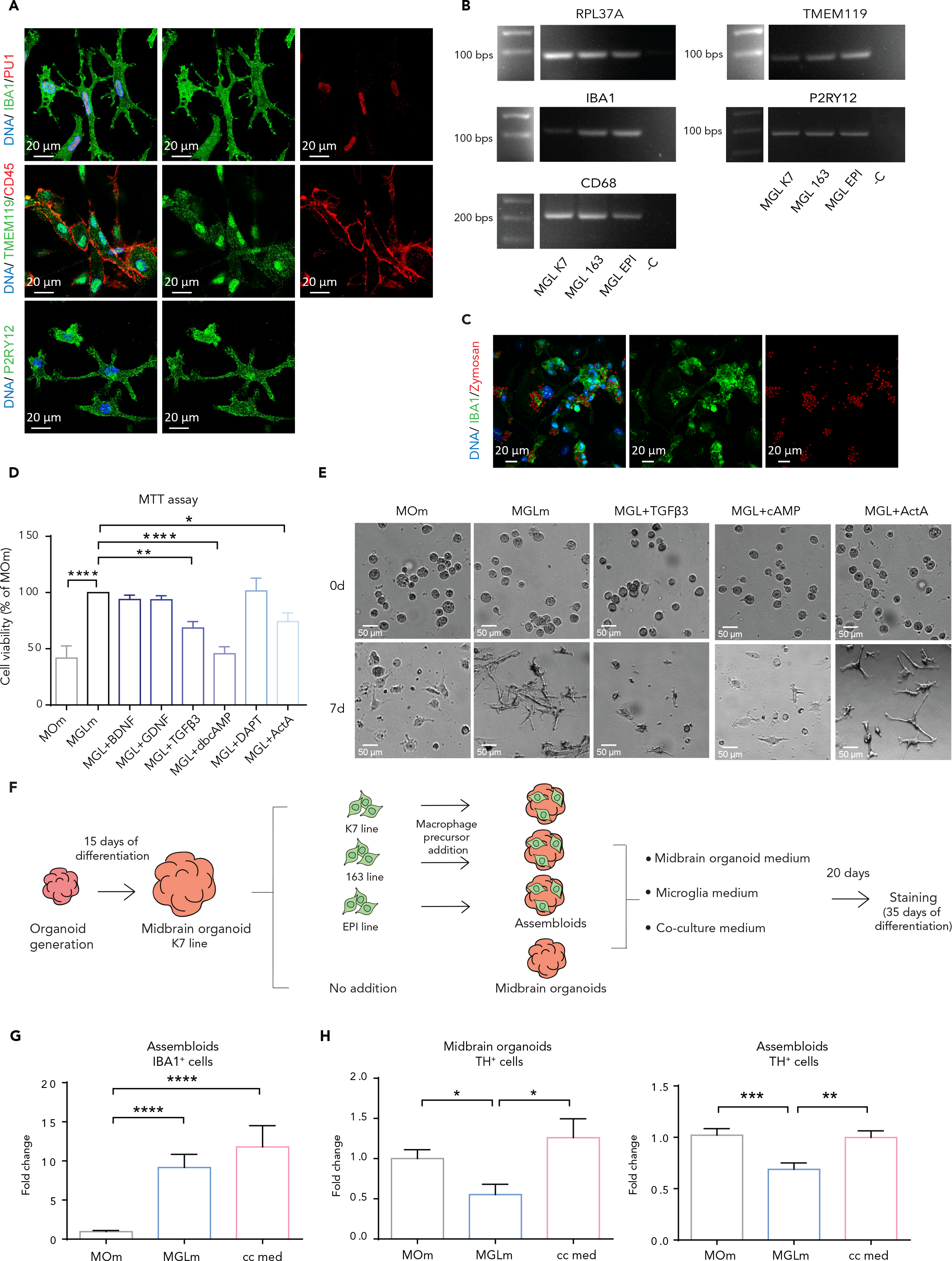
iPSC-derived microglia express specific markers, have phagocytosis ability and are compatible with the engineered co-culture medium. A. Immunofluorescence staining of microglia from line 163 for IBA1, PU.1 (upper panels), TMEM119 and CD45 (middle panels) and P2RY12 (bottom panels). B. IBA1, CD68, TMEM119 and P2RY12 gene expression in microglia from the three used lines. The Ribosomal Protein RPL37A coding gene was used as a housekeeping gene due to its stable expression. C. Immunofluorescence staining of microglia from the EPI line for IBA1 and Zymosan. For lines K7 and 163 see Figure S1B. D. Cell viability of 2D microglia from lines 163 and EPI (MTT assay) after 10 days of treatment with midbrain organoid media (MOm) or microglia medium (MGLm) without further supplementation or supplemented with neurotrophic factors. (n=3, 3 batches). E. Representative bright field images of the microglia morphology (line 163) at day 0 and 7 of culture with MOm, MGLm, MGLm supplemented with TGF β3 (MGL + TGF β 3), with cAMP (MGL + cAMP) or with Activin A (MGL + ActA). F. Schematic diagram of the steps for the media optimization in assembloids and midbrain organoids. G. IBA1 positive (IBA1^+^) population in assembloids upon culture with MOm, MGLm or co-culture medium (cc med). Y axis represents the fold change with respect to the control (MOm). H. TH positive (TH^+^) neuron population in midbrain organoids (left) and assembloids (right). Y axis represents the fold change with respect to the control (MOm) (n (midbrain organoids) = 2, 2 batches, n (assembloids) = 5, 2 batches and 3 cell lines). Data are represented as mean ± SEM. *p<0.05, **p<0.01, ***p<0.001, ****p<0.0001 using a 2way ANOVA with Dunnett’s multiple comparisons test. Abbreviations: BDNF, brain-derived neurotrophic factor; GDNF, glial cell-derived neurotrophic factor; TGFβ3, Transforming growth factor beta-3, cAMP, cyclic adenosine monophosphate; ActA, activin A.

A major challenge in assembloid generation is the compatibility of cell culture conditions for different cell types. In order to assess the microglia compatibility with the organoid culture medium, we first examined the toxicity of each midbrain organoid medium supplement on microglia survival. Macrophage precursors were cultured for 10 days with organoid maturation medium (containing BDNF, GDNF, TGFβ3, db cAMP, DAPT and Activin A), microglia differentiation medium (containing IL-34 and GM-CSF), or microglia medium individually supplemented with each of the neurotrophic factors contained in the organoid medium. After 10 days of exposure to the different media, an MTT viability assay showed a significant decrease in cell viability in the presence of TGFβ 3 (69.18% ± 8.371, p=0.0024), db cAMP (47.10% ± 8.371, p<0.0001) and Activin A (76.01% ± 8.371, p=0.0292) compared to microglia medium (Figure 1D and E). After only 7 days of exposure, an impairment of differentiation and a lower viability were already observed visually in the cells cultured with organoid medium as well as the microglia medium supplemented with TGFβ3 or db cAMP (Figure 1E). These results indicate that the midbrain organoid medium, containing TGFβ 3, db cAMP and Activin A might not be suitable for the cultivation of microglia containing midbrain organoids.

### The co-culture medium allows microglial survival and neuronal differentiation in midbrain organoids

After testing the organoid medium on microglia, we next assessed the effects of the different media compositions on the neuronal population of the midbrain organoids. Taking into consideration the cytotoxic effect on 2D monoculture microglia, we combined the microglia differentiation medium with the least microglia-toxic neurotrophic factors from the organoid maturation medium. Hence, we supplemented the microglia differentiation medium with BDNF, GDNF, DAPT and Activin A – since they play an important role in neuronal maturation - and refer to this medium combination as the “co-culture medium” (Table S2). In order to study the effects of the co-culture medium on the organoids, we used midbrain organoids from the line K7 as a control, and 3 groups of assembloids: K7 organoids co-cultured with K7 macrophage precursors, K7 organoids co-cultured with 163 macrophage precursors and K7 organoids co-cultured with EPI macrophage precursors (Figure 1F. The “assembloids” group in graphs represents the pooled results from the three co-cultured groups). We cultured those groups with midbrain organoid medium, microglia differentiation medium or co-culture medium for 20 more days (Figure 1F). As expected, the IBA1 positive microglial population was significantly reduced when organoid medium was used, whereas no difference was observed when cultured with microglia or co-culture medium (Figure 1G, Figure S1C). The major neuronal cell type in midbrain organoids are TH positive dopaminergic neurons. Interestingly, we observed a significant decrease in TH positive neurons in both midbrain organoids and assembloids in the presence of microglia differentiation medium (Figure 1H, Figure S1C). This decrease was not observed in the MAP2 positive pan neuronal population (Figure S1D, Figure 2C, Figure S1I). The reduction of TH positive cells indicates an impairment of dopaminergic neuron differentiation under microglia medium culture conditions. Notably, in the presence of co-culture medium, the levels of TH in both midbrain organoids and assembloids remained at similar levels as for the organoids cultured with midbrain organoid maturation medium (Figure 1H, Figure S1C). In summary, we here developed a unique co-culture medium that is optimal for both microglial survival and dopaminergic neuron differentiation in human midbrain organoids.

**Figure 2.**
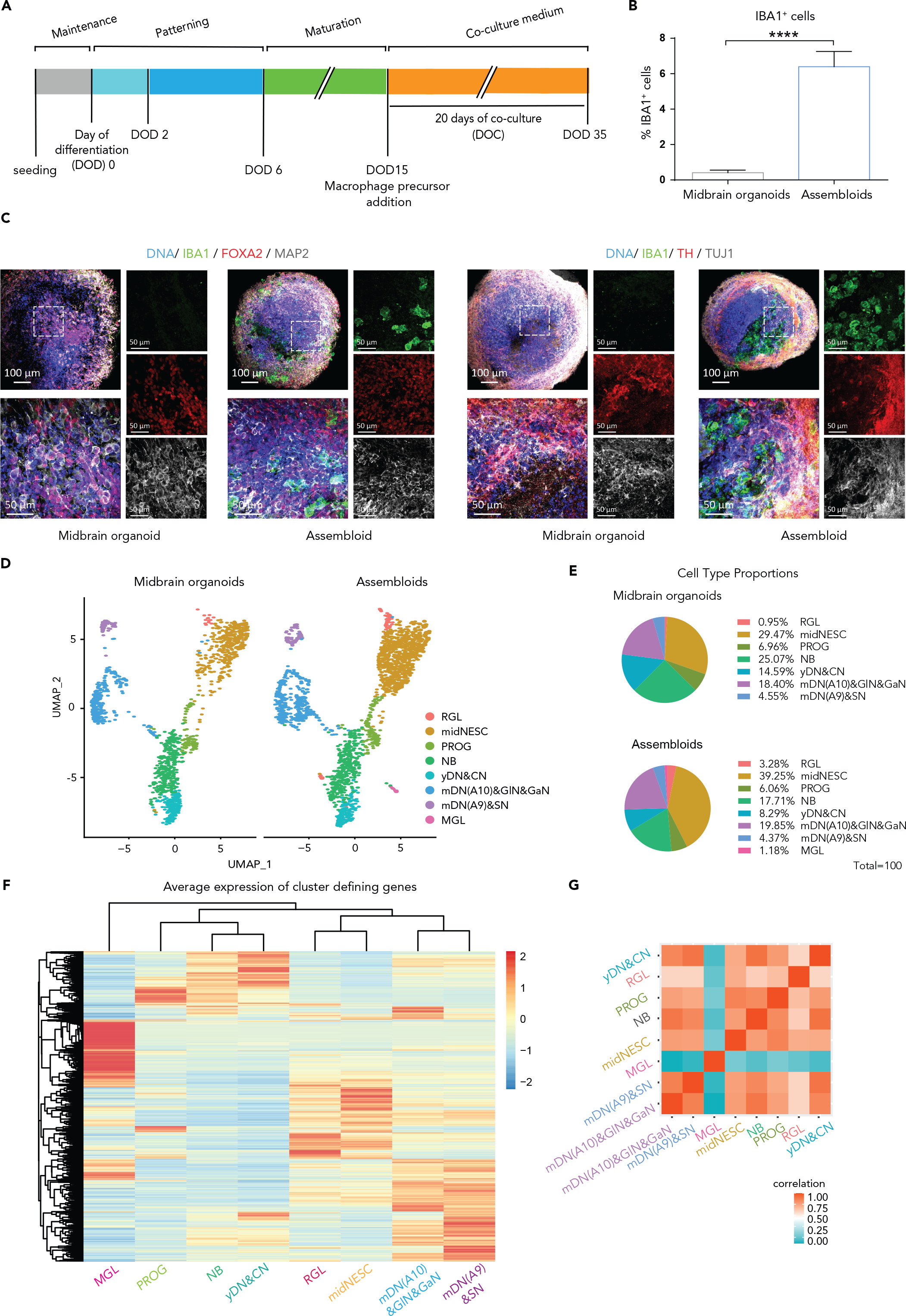
The co-culture medium allows a successful microglia integration and seven other neural cell populations in assembloids. **A.** Timeline of the co-culture of midbrain organoids with macrophage precursors. DOD = Day of Differentiation, DOC = Day of co-culture. **B.** IBA1 positive (IBA1^+^) cell percentage in midbrain organoids and assembloids. Assembloids present around 6.4% of IBA1^+^ cells (n (midbrain organoids) = 5, 5 batches, n (assembloids) = 15, 5 batches, 3 cell lines). Data are represented as mean ± SEM. **C.** Immunofluorescence staining of midbrain organoids and assembloids with microglia from the line K7 for IBA1, FOXA2 and MAP2 (left panels), and for TH and TUJ1 (right panels). For lines 163 and EPI see Figure S1H. D. UMAP visualization of scRNA-seq data - split by microglial presence - shows 8 different defined cell clusters in assembloids. RGL, radial glia; midNESC, midbrain specific neural epithelial stem cells; PROG, neuronal progenitors; NB, neuroblasts; yDN&CN, young dopaminergic and cholinergic neurons; mDN(A10)&gaN&GlN, mature A10 specific dopaminergic neurons, gabaergic and glutamatergic neurons; mDN(A9)&SN, mature A9 specific dopaminergic neurons and serotonergic neurons, MGL, microglia. **E.** Proportions of different cell types in midbrain organoids and assembloids**. F.** Average expression of cluster defining genes in assembloids. **G.** Spearman’s correlation between different cell types in assembloids. *p < 0.05, **p<0.01, ***p<0.001, ****p<0.0001 using a Kruskal-Wallis 1way ANOVA with Dunn’s multiple comparisons test (for MGLm and cc med vs MOm), and a Mann-Whitney test (for MGLm vs cc med) in A and B, and a Mann-Whitney test in E.

### Assembloids contain IBA1 positive functional microglia and express midbrain specific neuronal markers

After successfully optimizing the co-culture conditions, midbrain organoids were generated using a pure ventral midbrain-patterned NESC population (Figure S1E) from the hiPSC line K7 (Smits *et al*., 2019; Nickels *et al*., 2020). After 2 days of maintenance, and 15 days of dopaminergic neuron differentiation induction, we proceeded to the co-cultures (Figure 2A).

They were performed as described in the previous section, using K7 midbrain organoids as controls, and co culturing K7 organoids with K7 macrophage precursors, with 163 macrophage precursors or with EPI macrophage precursors (following Figure 1F excluding the media testing). During the whole study, we did not observe any major line specific differences. During the co-culture, the macrophage precursors first aggregated, forming smaller round colonies that attached to the surface of the organoid and then incorporated within that same structure (Figure S1F).

After 20 days of co-culture (35 days of dopaminergic differentiation), we validated the incorporation of microglia into the organoids by IBA1 immunostaining. We then assessed the percentage of IBA1 positive cells in assembloids using a previously published computer-assisted image analysis pipeline for marker identification and quantification with small modifications (Smits *et al*., 2019). On average, 6.4% of the assembloid cells were IBA1 positive (Figure 2B). The morphology of the incorporated microglial cells varied between round and partially ramified (Figure S1G). We confirmed the presence of a neuronal population by a Beta-tubulin III (TUJ1) staining, and assessed further neuronal differentiation by MAP2 staining. In addition, FOXA2 positive midbrain specific dopaminergic neuron precursors and TH positive differentiated dopaminergic neurons were observed in assembloids (Figure 2C, Figure S1I). After culturing the assembloids for 70 days, we further confirm the presence of a GFAP positive astrocytic population in the assembloids (Figure S1H).

### Single nuclear RNA-sequencing reveals eight different cell populations within midbrain- microglia assembloids

In order to further characterize the microglia within assembloids we performed single nuclear RNA-sequencing (snRNA-Seq). Separation of clusters in cells coming from midbrain organoids and assembloids clearly showed that the microglia cluster is specific for assembloids and not present in midbrain organoids (Figure 2D). Cell types were identified based on marker gene expression from La Mano et al. 2016 (G *et al*., 2016). Moreover, the identified cell clusters were validated and neuronal subtypes were specified (Figure S2 and Table S5). Quality controlled clustering of single cells (Figure S3) showed eight different cell types: radial glia (RGL), midbrain specific neural epithelial stem cells (midNESC), neuronal progenitors (PROG), neuroblasts (NB), young dopaminergic and cholinergic neurons yDN&CN, mature A10 specific dopaminergic neurons, GABAergic and glutamatergic neurons (mDN(A10)&GABA&GLUT), mature A9 specific dopaminergic neurons and serotonergic neurons (mDN(A9)&SN) as well as microglia (MGL, Figure S2).

Proportions of the different clusters in midbrain organoids and assembloids are represented in pie charts (Figure 2E). midNESC were the most prominent cell type by representing 35% of total cells. More than 60% of the organoid and assembloid cells showed neuronal identity (corresponding to the groups NB, yDN&CN, mDN(A10)&GlN&Gan, mDN(A9)&SN). Less than 5% were represented by glial cells (astrocytes, oligodendrocytes and microglia) at this stage of differentiation. Because of a considerable sample and cell loss, we believe the proportion of microglia cells represented in the Figure 2E is not accurate. However, we proceeded with the analysis of microglial identity and functionality-related genes, as well as for the other cell clusters. Progenitors were closest to neuroblasts and young neurons, whereas radial glial cells were very similar to midNESCs. The different mature neuronal groups clustered together. Clustering of the marker genes showed the relation between different cell clusters (Figure 2F). As expected, MGL cells clustered apart from all other cell types. A distinct genetic signature of the microglia cells was further validated by a Spearman correlation analysis (Figure 2G).

### Microglia in assembloids show a typical immune cell signature

Next, we aimed at identifying gene signatures unique to microglia in comparison to the other cell types in assembloids. Firstly, we used the expression of the 100 most variable genes obtained from the snRNA-Seq analysis to cluster the different cell populations observed in assembloids (Figure 3A). Indeed most variability between cells clusters came from the microglia, which had a completely different genetic signature (Figure 3A). Then, we identified the top 100 markers defining microglia identity (Figure 3B). From those, several canonical marker genes were chosen in order to characterize the main functions of microglia. General microglia marker genes such as *IBA1* and *PU1* were significantly and specifically expressed in microglia. In addition, the microglia core signature revealed genes involved in antigen presentation (*HLA-DMB*), cytokine and complement signaling (*CSF1R, IL18, C1QC*) as well as pathogen and self-recognition such as toll-like receptor signaling (*TLR2*), C-type lectins (*CLEC7A*) and mannose and nod-like receptors (*MRC1, NAIP*). Moreover, genes involved in microglial adhesion (*ITGAM*) and motility through chemokine signaling (*CCL2*), as well as purinergic signaling (*P2RX4*) were significantly expressed in microglia (Figure 3C). Furthermore, several chemokines and cytokines were expressed predominantly in microglia (Figure 3D).

**Figure 3.**
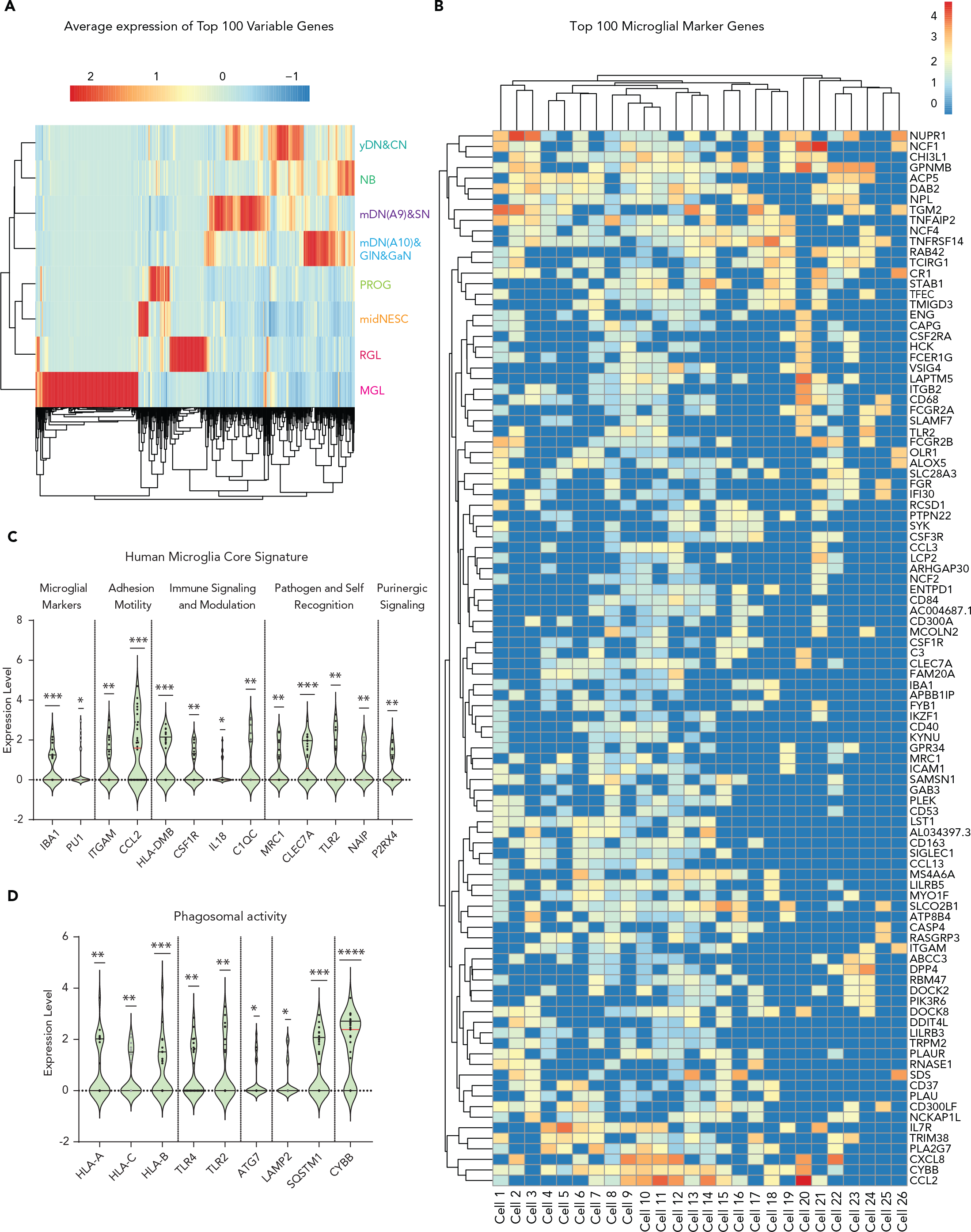
Microglia in assembloids have a typical immune cell signature. **A.** Heatmap showing the average expression of 100 most variable genes between midbrain organoids and assembloids. **B.** Expression of top 100 microglial marker genes across microglia cells in assembloids. **C.** Microglia core signature, expression of microglia marker genes as well as genes involved in adhesion, immune response, pathogen recognition and purinergic signaling. **D.** Gene expression levels of genes related to phagocytic activity. Violin plot shows average expression level. *p <0.05, **p<0.01, ***p<0.001, ****p<0.0001 using a Wilcox one-sample tests, median in red, quantiles in black. Dots represent single cells.

In order to address microglia functionality, we then measured chemokine and cytokine release in midbrain organoids and assembloids. For that, culture medium from midbrain organoids and assembloids at day 20 of co-culture was used. The medium collected was in contact with the organoids or assembloids for 3 days. As expected, in a hierarchical clustering analysis, midbrain organoids clustered in a different group than the microglia containing assembloids (Figure 4A). For a total of 18 analyzed cytokines (IL-7, IL-12p17, IL-3, TNFα, IL-1α, IL-1, IL-6, IFNα, IL-

**Figure 4.**
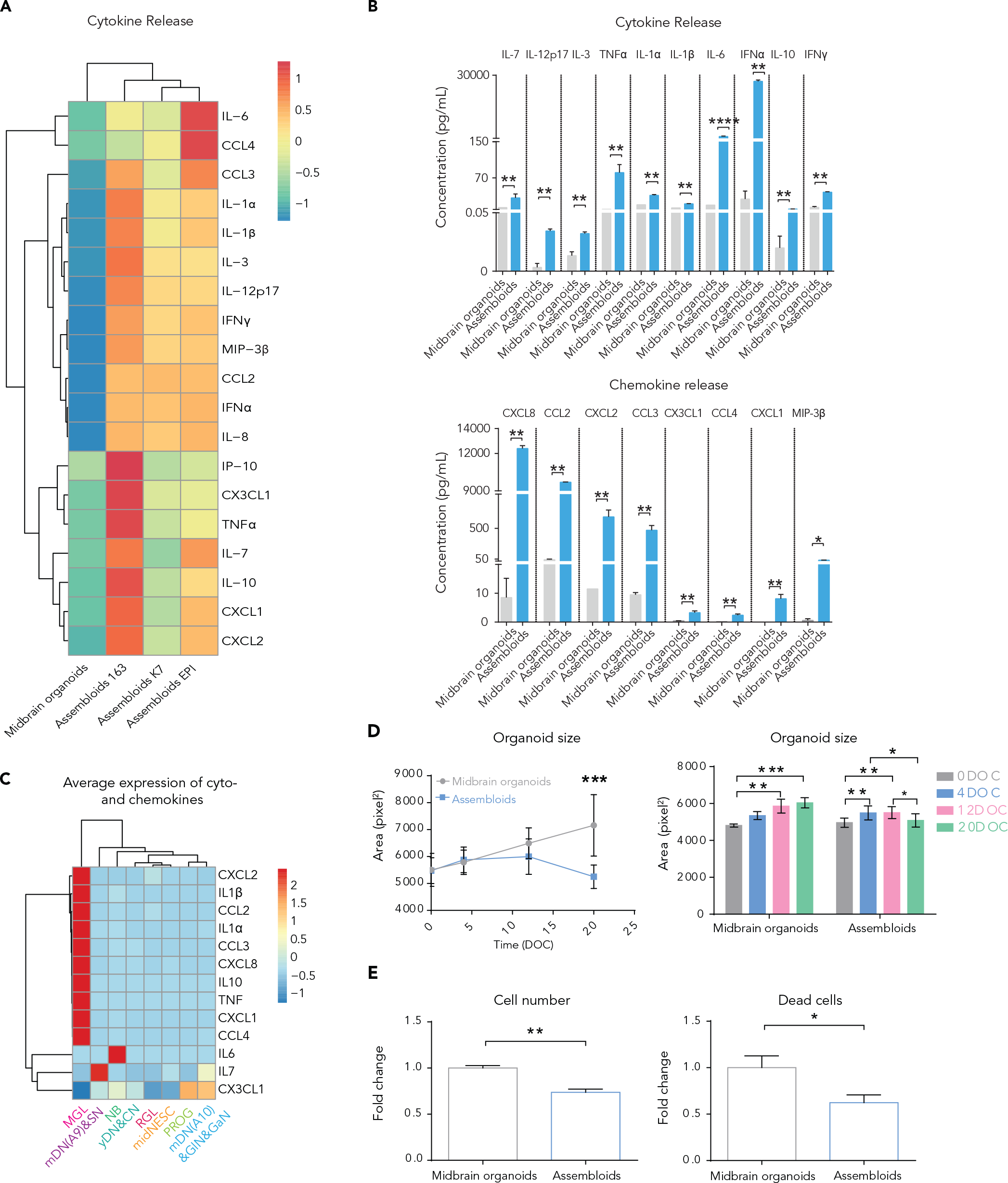
Microglia in assembloids have phagocytic capacities and release cytokines and chemokines. **A.** Heatmap representing the measured levels of cytokines and chemokines in cell culture media (pg/ml) shows two different clusters corresponding to midbrain organoids and assembloids (n=9, 3 batches, 3 lines). **B.** Cytokine (upper graph) and chemokine (bottom graph) levels in midbrain organoids and assembloids (n=9, 3 batches, 3 lines). **C.** Average expression of cytokine and chemokine genes across cell types. **D.** Organoid surface area in midbrain organoids and assembloids over time (left graph, n=3, 3 batches). Comparison of the organoid size, measuring the same organoids and assembloids, in four time points during culture (right graph, n=3, 3 batches). **E.** Cell number (total nuclei count, left panel) and dead cells (pyknotic nuclei, right panel) in midbrain organoids and assembloids after 20 days of culture (n (midbrain organoids) = 5, 5 batches, n (assembloids) =15, 5 batches, 3 lines). Data are represented as mean ± SEM. *p < 0.05, **p<0.01, ***p<0.001, ****p<0.0001 using a Mann-Whitney test in A and B, a multiple t test with the Holm-Sidak method for D (left panel), a 2way ANOVA with Tukey’s multiple comparisons test for D (right panel) and a Mann-Whitney test in E.

10 and IFNγ) and chemokines (CXCL8, CCL2, CXCL2, CCL3, CXCL1, CCL4, CX3CL1 and MIP-3 ) we observed a significant increase in the medium from assembloids compared to midbrain organoids, suggesting that immune functionality was acquired through the microglia incorporation (Figure 4B). Interestingly, although the RNA expression levels for IL-6, IL-7 and CX3CL1 were higher in non-microglia cells (Figure 4C), the levels of released cytokines were higher in the assembloids medium (Figure 4B). Hence, potentially the released cytokines are not necessarily produced in the microglia, but their secretion is stimulated by the presence of microglia.

### Microglia in assembloids are functional and affect the expression of cell survival-related genes

As we previously demonstrated, microglia have the ability to phagocytose Zymosan particles in 2D monoculture (Figure 1C). Therefore, we assessed if microglia in assembloids express genes involved in phagocytosis. Indeed, in assembloids we observed a higher expression of phagocytic genes suggesting the presence of a functional phagolysosomal pathway (Figure 3D). Genes included antigen recognition (*HLA-A,B,C*), receptor (*TLR2,4*) and internal signaling (*ATG7*), as well as autophagic vesicle formation (*LAMP2*, *SQSTM1* (*p62*)) and lysosomal degradation/oxidation (*CYBB*). These results indicate these microglia should have phagocytic capacity and hence the ability to remove dead cells. Interestingly and in agreement with this hypothesis, when we measured the organoid size throughout the culture period we observed a decrease of the assembloid size over time (Figure 4D). Midbrain organoids and assembloids were stained using the DNA dye Hoechst, and using our computer-assisted image analysis pipeline for cell type segmentation(Smits *et al*., 2019), we quantified Hoechst signal. Levels of total cells, live cells and necrotic/late apoptotic cells were evaluated. We observed that the total amount of cells was lower in assembloids compared to midbrain organoids, correlating with the size measurements. Particularly, the amount of dead cells was significantly lower in the assembloids (Figure 4E), suggesting that microglia may eliminate dead cells in assembloids.

### Microglia have an effect on oxidative stress and immune response in assembloids

After establishing the successful integration of microglia into midbrain organoids and showing important aspects of microglia functionality, we investigated the potential influence of microglia on the neural cells in midbrain organoids. We performed differential gene expression analysis over the neural cells in assembloids and midbrain organoids, which revealed 423 significantly different genes (p<0.05). The top 100 differentially expressed genes (DEGs) across all cells are represented in a heat-map and clustered by cell type (Figure S4A). We assigned DEGs to each cell type and represented their overlap in Venn diagrams (p<0.05, Figure S4B). The left Venn diagram shows an overlap of all three neuronal clusters and the right Venn diagram overlaps NB, RGL, PROG and midNESCs. Most DEG were cell type specific, although some cell types shared DEGs, which was especially the case between both mature neuronal clusters.

Next, a pathway enrichment analysis was performed MetaCore using the DEG across all cell types in midbrain organoids and assembloids (Figure 5A). The analysis revealed 12 significantly enriched biological pathways. Besides ribosomal and cytoskeletal genes, genes for oxidative stress (10/160 genes), the immune response (12/242 genes) as well as neurogenesis and axonal guidance were significantly different (13/229). Interestingly, we found 10 enriched genes involved in hypoxia and oxidative stress. Among those, six genes showed a clear downregulation of this pathway in the presence of microglial cells; the expression of mitochondrial cytochrome oxidase 1 (complex IV) (*COX1*), peroxiredoxin1 (*PRDX1*), superoxide dismutase 1 (*SOD1*), glutathione peroxidase 4 (*GPX4*), ATPase (complex V) (*MT-ATP6*), as well as glutathione S-transferase 1 (*GSTP1*) was significantly lower in assembloids (Figure 5B).

**Figure 5.**
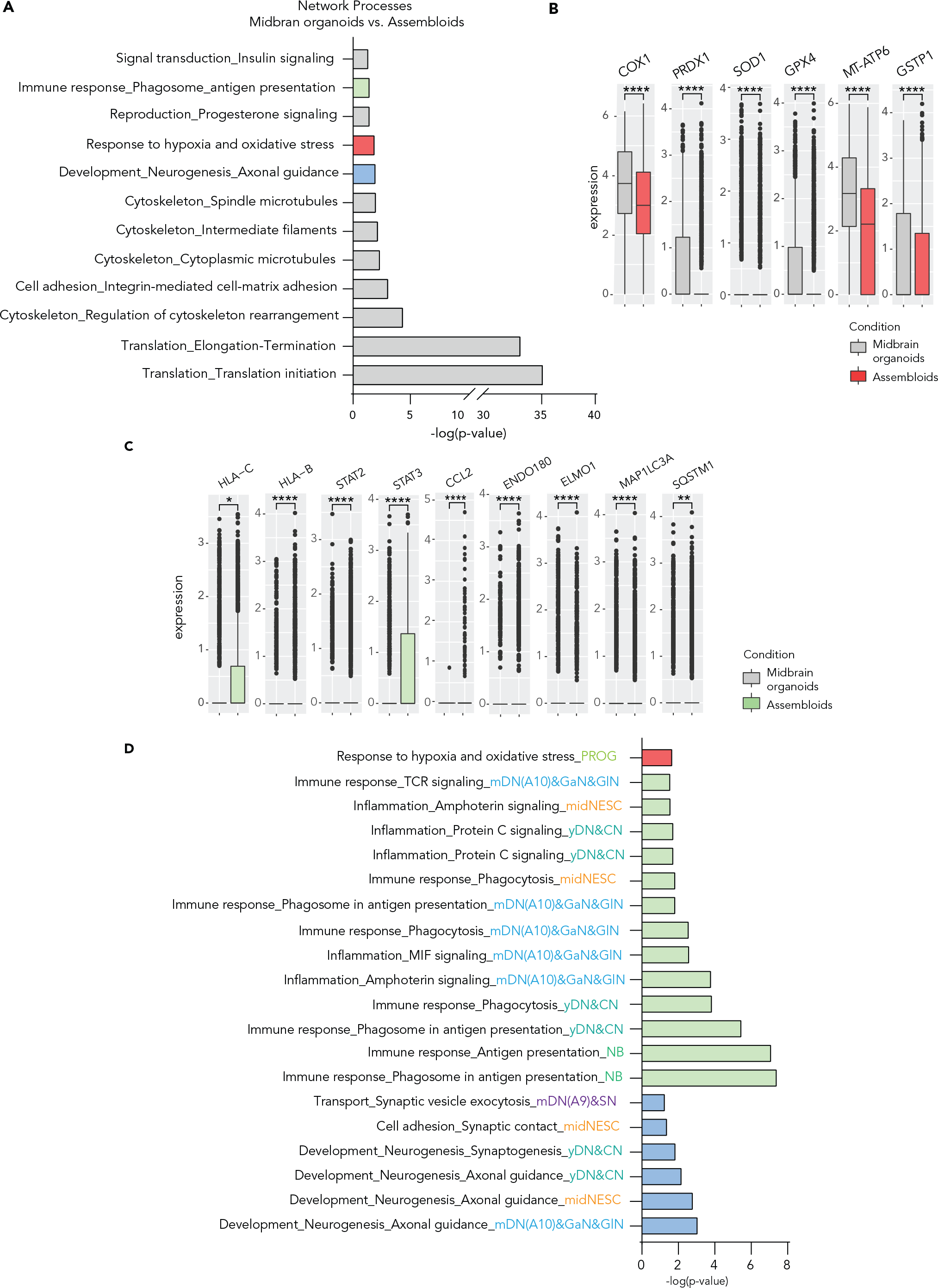
Microglia affect the expression of oxidative stress and immune response-related genes in assembloids. **A**. Differentially expressed gene enrichment analysis in assembloids against midbrain organoids reveals 12 significant network process pathways (FDR<0.05). **B.** Expression of genes related to response to oxidative stress in assembloids compared to midbrain organoids. **C.** Gene expression of genes related to immune response of assembloids. The presence of microglia increases the expression of genes related to antigen presentation and immune response, and decreases the expression of those related to autophagy in non-microglia cells. Box plots show mean expression and standard deviation. **D.** Enrichment analysis of cluster specific DEG p<0.05 between assembloids and midbrain organoids reveals significant FDR<0.05 network processes involved in oxidative stress, immune response as well as synaptic regulation. Dots represent single cells. Data are represented as mean ± SD. *p < 0.05, **p<0.01, ***p<0.001, ****p<0.0001 using a Wilcox test.

Next, we assessed the 12 genes involved in the immune response and in the antigen presentation processes (Figure 5C). Indeed, non-microglial cells expressed genes from the MHCII such as *HLA-C* and *B*. Moreover, the *STAT2* and *STAT3* genes were upregulated in the presence of microglia. The MRC2 (ENDO180) receptor involved in collagen internalization and remodeling was upregulated, while genes involved in cytokine mediated phagocytosis (*ELMO1*) and autophagy (MAP1LC3A (LC3), SQSTM1 (p62)) were downregulated (Figure 5C). An enrichment analysis of the DEGs for each cell cluster showed differences in genes involved in immune response, inflammation, phagocytosis and response to hypoxia and oxidative stress, among others (Figure S4C).

Microglia are important to main brain homeostasis. However, neuro-inflammation might occur when this homeostasis is compromised. Therefore, we assessed the expression of genes involved in pyroptosis – inflammation related cell death through inflammasome activation – including CASP1, NLRP3, PYCARD (data not shown) PPIA (Figure S5A). Neuro-inflammation related genes were unchanged or decreased in assembloids, suggesting that presence of microglia in organoids does not lead to neuro-inflammation in physiological conditions.

### Microglia affect the expression of synaptic remodeling-related genes in assembloids

Enrichment analysis of the DEGs for each cell cluster showed differences in genes involved in synaptic vesicle exocytosis, synaptic contact, synaptogenesis and axonal guidance in assembloids (Figure 5D and Figure S4C). To further investigate the microglia effects on synaptic pathways, we performed an extensive analysis of genes involved in synaptic processes. In assembloids, general synaptic markers such as *Synaptotagmin* (*SYT1*) and *Synaptophysin* (*SYP*), as well as the dopaminergic neuron circuit formation genes *ROBO1* and *DCC* were significantly downregulated across cell types (Figure 6A). Other important genes involved in synaptic vesicle exocytosis – such as *VMAT2* and *SNAP25* - were differentially expressed (Figure S5B). Moreover, we assessed axonal guidance and growth molecules, such as semaphorins (*SEMA3C*, *DPYSL2*), plexins, ephrins (*EPHA5*), neuropilins, neurofilaments and actin cytoskeleton (*NEFM, ACTB*) (Figure S5C). These genes were all differentially expressed, depending on the cell type, indicating that axonal remodeling is influenced by the presence of microglia. Furthermore, we examined some cell specific genes in mature neurons involved in action potential (*CASK, CACNA1A, CACNA1E, CACNA1B*) and active zones (*HCN1, KCNC3, KCND3*) within synapses and observed that those genes are upregulated in assembloids (Figure 6B). Apart from assessing the synapse-related gene expression, we observed reduced protein levels of the synaptic vesicle protein VAMP2 in assembloids compared to midbrain organoids (Figure 6C). Together these results suggest a role of microglia in synaptic remodeling and maturation within midbrain organoids.

**Figure 6.**
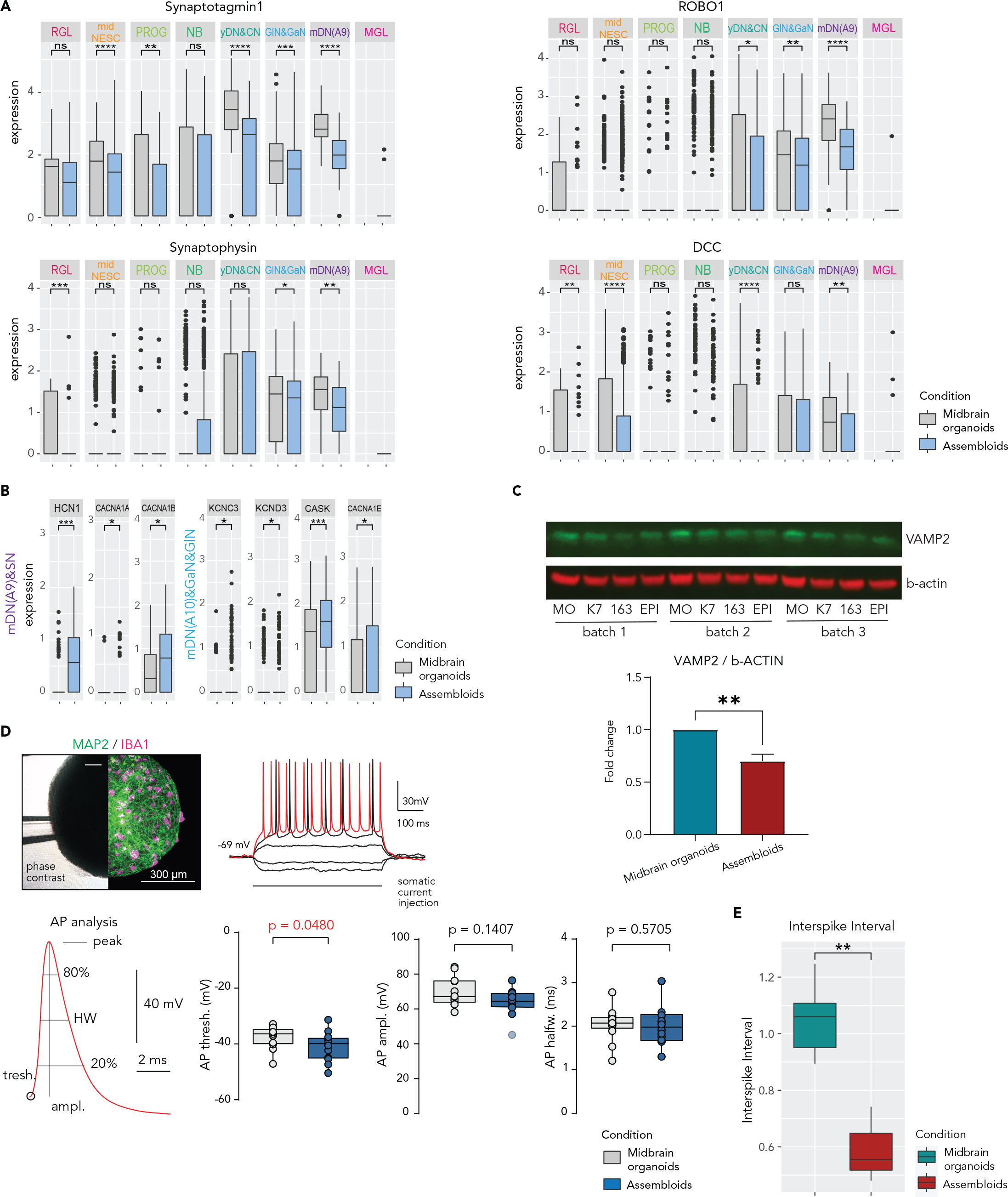
Microglia affect the expression of genes related to synaptic remodeling in assembloids and develop mature electrophysiological characteristics. A. Gene expression of general synaptic markers such as *Synaptotagmin* (*SYT1*) and *Synaptophysin* (*SYP*), and the dopaminergic neuron circuit formation genes *ROBO1* and DCC across cell clusters in midbrain organoids and assembloids. Data are represented as mean ± SD. * p< 0.05 using a Wilcox test. Dots represent single cells B. Expression of genes involved in action potential (*CASK, CACNA1A, CACNA1E, CACNA1B*) and active zones (*HCN1, KCNC3, KCND3*) within synapses in the dopaminergic neuron cluster (mDN(A9)&SN) in midbrain organoids and assembloids. Data are represented as mean ± SD. * p< 0.05 using a Wilcox test. Dots represent single cells. C. Western blot showing protein levels of the synaptic vesicle marker VAMP2 and the β-actin (upper panels). Bar graph showing the Western blot quantification from the upper panels (n (midbrain organoids) = 3, 3 batches, n (assembloids) = 9, 3 batches, 3 cell lines). **p<0.01 using a Mann-Whitney test. Data are represented as mean ± SEM. D. Fixation of organoid during recording and post hoc verification of microglia presence. The left half shows an infra-red phase-contrast life image with the fixation pipette (upper left panel). The right half shows the same organoid after immunofluorescence staining for MAP2 and IBA1. Example traces show voltage response to hyperpolarizing and depolarizing current injections of a neuron inside an assembloid measured by whole-cell patch-clamp. Voltage responses that exhibited action potential (AP) following 50ms after stimulus onset were used for AP analysis (upper right panel). Analysis of AP waveform characteristics (bottom left panel). Voltage thresholds were significantly more depolarized in assembloid neurons (bottom middle-left panel, n =14 neurons in midbrain organoids and n =13 cells in assembloids), although analysis of AP amplitude (bottom middle-right panel) and half width (bottom right panel) shows no systematic differences between both groups. Box plots indicate median, 25th, and 75th percentiles and raw data points. Outliers deviating 2.5 standard deviations are marked translucent and were excluded from statistical analysis for normally distributed data. P-values were determined using unpaired t-tests or Mann-Whitney rank test (indicated as p, see Supplemental Experimental Procedures). D. Multi-electrode array results showed a lower inter-spike interval in assembloids compared to midbrain organoids (n (midbrain organoids) = 3, 3 batches, n (assembloids) = 9, 3 batches, 3 cell lines). ***p=0.001, **p=0.01, *p=0.05 using a Wilcox test. Boxes cover data from the first to the third quartile. The whiskers go from each quartile to the minimum or maximum.

In order to investigate the functional impact of microglia in midbrain organoids we performed electrophysiological measurements of passive and active membrane properties as well as firing behavior. We performed patch-clamp experiments of visually identified neurons in the intact organoids from 20 to 35 days after microglia addition (Figure 6D). Neurons in midbrain organoids and assembloids exhibited similar resting membrane potentials and input resistances and reliably fired repetitive action potentials in response to somatic current injections (n midbrain organoids =14; n assembloids = 13 cells; Figure S5D). Depolarizing steps in voltage-clamp configuration triggered strong inward currents in all tested neurons, indicative of fast voltage-activated sodium currents (Figure S5E). The amplitude of these currents was not different between both groups. Firing characteristics and action potential waveforms (Figure 6D) varied considerably, which was expected from neurons at different degrees of maturation.

Importantly, the voltage threshold for the action potential generation was more negative in the group of assembloid neurons (-39.86 (midbrain organoids) against -36.35 (assembloids) ± 3.281, p=0.0480, Fig 6d), which is a common and strong indicator of increased neuronal excitability in mature neurons. In sum, neurons in assembloids develop fully mature electrophysiological properties with a lower threshold for action potential generation than in midbrain organoids. Furthermore, we performed multi-electrode array (MEA) analysis with midbrain organoids and assembloids containing microglia from line K7, 163 or EPI (Figure 6E). The results showed that, after 40 days of co-culture, which corresponds with the time point when the Patch clamp measurements were performed, assembloids had a lower interspike interval than midbrain organoids (Figure 6E). These results back up the observed lower action potential threshold observed in the patch-clamp experiment.

In order to support these findings and investigate further the differences between midbrain organoids and assembloids, we performed a non-polar exo-metabolomic analysis from culture supernatants 20 days after microglia addition. The assay showed a different metabolic profile in midbrain organoids compared to assembloids (Figure S5F). A total number of 14 metabolites were significantly different. Among them, we observed a higher uptake of glucose and pyruvic acid from assembloids (Figure S6A). Regarding amino-acid metabolism, we observed a lower secretion of phenylalanine, tyrosine, methionine, lysine, putrescine, threonine, leucine, isoleucine and valine by the assembloids compared to midbrain organoids. Furthermore, the levels of uptaken asparagine and serine from the medium were higher in assembloids. The secretion of glutamine was higher assembloids (Figure S6B).

## Discussion

### Microglia can be efficiently integrated into Midbrain organoids

Here we described the generation of human midbrain assembloids containing microglia as well as the effect that microglia have on midbrain organoid structure and function. Although some studies have shown a microglia-like population in brain organoids (Mansour *et al*., 2018; Ormel *et al*., 2018), integration of human iPSC-microglia cells into midbrain organoids has not previously been reported. Moreover, particularly the assessment of how microglia are affecting brain organoid physiology is heavily under-investigated.

In this study, we have shown for the first time the successful incorporation of a significant amount of microglia into midbrain organoids, obtaining an average of 6.4% of IBA1 positive cells in assembloids. Studies based on immunocytochemistry show around 10% of IBA1 positive cells in human substantia nigra (Mittelbronn *et al*., 2001). One of the main challenges we faced during the integration were the cytotoxic effects of the midbrain organoid medium on microglia, and the unsuitability of the microglia differentiation medium on the organoids. Interestingly, while TGF-beta 1 and 2 have been associated with microglia function and inflammatory response (Hu *et al*., 1995; Lieb, Engels and Fiebich, 2003; WK *et al*., 2004; Taylor *et al*., 2017) , very little is known about the relationship between TGF-beta 3 and microglia. TGF-beta 3 has been associated with dopaminergic neuron differentiation (Roussa *et al*., 2006), which is the reason why the dopaminergic neuron differentiation medium contains this recombinant molecule. However, no studies associating a beneficial relationship between TGF-beta3 and microglia have been found. We hypothesize that this subtype of the TFG-beta family may be acting differently than TGF-beta 1 and 2, since the 3D structure of TGF-beta 1 and 2 are considerably different than TGF-beta3, which could lead to functional differences with respect to their effect towards microglia (Andrew P. Hinck *et al*., 1996; EV *et al*., 2000; Grütter *et al*., 2008). The absence of neurotrophic factors – which promote dopaminergic differentiation – in the microglia differentiation medium, led to a significantly lower amount of dopaminergic neurons in midbrain organoids and assembloids. This incompatibility might be one of the reasons why, so far, no studies have shown an efficient co-culture of dopaminergic neurons with microglia. Here, we developed an optimized co-culture medium, which allowed the survival of microglia within the organoids as well as an efficient dopaminergic neuron differentiation. The incorporated microglia clustered separately from the cells of ectodermal origin in the midbrain organoids, showing that both cell lines keep their cellular identity under co-culture conditions.

This is in line with in vivo development, as microglia are derived from primitive hematopoietic stem cells and as such have a completely different genetic signature (Alliot, Godin and Pessac, 1999; Ginhoux *et al*., 2010; Schulz *et al*., 2012). Unfortunately, and as stated before, the sample loss during the pre-processing for snRNA-Seq makes the cell proportions from the sequencing inaccurate. Thus, additional runs of sequencing would allow us to obtain more reliable results concerning the cell proportions. However, the obtained data is still of good quality to proceed with gene expression analysis for cell clustering and for assessing functionality-related genes.

Apart from clustering separately from the rest of the assembloid cells, microglial cells express genetic markers for all the major functions of the human microglial core signature (Galatro *et al*., 2017). This confirms that the population we integrated into the organoid is, indeed, microglial.

Until now, no studies analyzed which effect the integration of microglia cells have on the functionality of midbrain organoids. Our data demonstrate that the integration of microglia influence neural stress, cell death, neuronal cyto-architecture and synapse remodeling- related gene expression in the neuronal network.

### Microglia communicate with neurons and play a role in midbrain organoid stress response

Microglia have cell-cell communication ability through cytokine and chemokine signaling (Carbonell *et al*., 2005; Arnò *et al*., 2014; Siddiqui, Lively and Schlichter, 2016; Haenseler, Sansom, *et al*., 2017). They can move towards apoptotic cells in order to phagocyte and eliminate cell debris (Chan *et al*., 2003). Apoptotic neurons generate chemotactic signals recognized by microglial receptors, which attract them to the apoptotic area in order to phagocytose dying cells (Witting *et al*., 2000). Our data suggest that microglia are functional in assembloids. Phagocytic genes were significantly expressed. These gene expression data were experimentally confirmed through a 2D Zymosan phagocytosis assay. Furthermore, our gene expression analysis via snRNA-Seq showed the expression of multiple cytokines and chemokines, as well as cyto- and chemokine receptors, in microglia from assembloids. We also measured many cytokines and chemokines in culture media, and observed that their levels were significantly higher in assembloid medium compared to midbrain organoids medium. We observed a reduced size of assembloids compared to midbrain organoids, and a lower amount of dead cells in assembloids. These results lead us to hypothesize that microglia in assembloids may be attracted to apoptotic areas and phagocytose apoptotic cells and cell debris. High levels of cell death and absence of removal mechanisms for dead cells, particularly in the center of organoids, was previously a key limitation of organoid technology (Berger *et al*., 2018; Nickels *et al*., 2020).

Previous work on organoids suggested that neurons in organoids have unusual high stress levels (Bhaduri *et al*., 2020). Interestingly, the DEG analysis shows differences in oxidative cell stress-related genes (often linked to cell death). Moreover, the downregulation of autophagy within non-microglial cells might be linked to reduced overall stress due to starvation, which is in line with the increased metabolite uptake from the media. Additionally, the metabolomics analysis revealed a significantly lower amount of Leucine, Isoleucine, Valine, Phenylalanine and Tyrosine in the assembloid culture medium. Increased plasma levels of those metabolites have been associated with acute hypoxic exposure in rats (Muratsubaki and Yamaki, 2011).

We also observed upregulated genes involved in the immune response in assembloids. Non-microglial cells may have reacted towards microglia by upregulating antigen presenting factors, and STAT2 and STAT3 pathways involved in the inflammatory response. Although neuronal cells respond to microglia, no inflammasome activation was observed in the current study. This, together with reduced oxidative stress and cell death in assembloids, suggests that the communication could be neuroprotective. In addition, ER stress and UFPR remained unaffected or reduced in assembloids. Overall, these results indicates alterations in cell signaling, oxidative stress and inflammation. Further studies on cell death and oxidative stress would be of value to confirm the observed results and support our hypothesis.

### Microglia induce alterations in synaptic gene expression in assembloids

In the current study, the expression of synaptic marker genes is reduced in assembloids. Furthermore, protein levels of the pre-synaptic protein VAMP2 were lower in assembloids compared to midbrain organoids. Physiologically, microglia are responsible, among other functions, for remodeling the cyto-architecture and connectivity of the brain by stripping unnecessary or misguided synapses (Wake *et al*., 2009; Tremblay *et al*., 2011). Here we could show that genes in axonal guidance and cytoskeletal organization are deregulated upon microglia presence. Moreover, whereas the expression of most synaptic genes are reduced, the expression of genes involved in triggering action potential are increased. These results lead us to hypothesize that inactive synapses may be eliminated while active synapses are strengthened. This phenomenon would be in line with the previously described synaptic pruning function of microglia (Paolicelli *et al*., 2011; Sellgren *et al*., 2019). The metabolomics results may be supporting this theory; there was an increase in glucose metabolism in assembloids via a higher uptake of glucose and pyruvic acid from the medium. The higher metabolic activity in the glucose metabolism may indicate a higher need for substrates for the TCA cycle, which leads to amino-acid production and, eventually, to neuro-transmitter production (Tiwari, Ambadipudi and Patel, 2013). Further, we observed a lower release of methionine to the medium. Methionine is related to processes of neurotransmission and neuromodulation (Kurbat and Lelevich, 2009). The secretion of glutamine to the medium was higher in assembloids. Since this amino acid is directly linked to the neurotransmitter glutamate production, this might indicate a higher production of glutamate. The uptake of the amino acids phenylalanine and tyrosine was higher in assembloids. Interestingly, tyrosine can be metabolized from phenylalanine, which can be metabolized to L-DOPA and, after that, to dopamine (Weinberg *et al*., 2019). Further experiments assessing synapse pruning would be of great value to confirm these observations.

Finally, the electrophysiology analysis showed that neurons in assembloids form a functional network. Interestingly, the voltage threshold for action potential generation, a common marker of neuronal excitability, was lower in assembloids compared to midbrain. Furthermore, the MEA results show a lower inter-spike interval in assembloids compared to midbrain organoids. Together with a tendency to higher membrane potentials, this indicates elevated levels of excitability in these neurons. Since the intrinsic excitability of neurons is often fine-tuned to the amount and frequency of external inputs and network activity, this change might in fact compensate for the possible degree of synaptic pruning previously discussed. Furthermore, these results are in line with previous reports, which describe that microglia increase neuronal excitability (Klapal, Igelhorst and Dietzel-Meyer, 2016).

Thus, our results suggest that microglia in assembloids may perform synaptic pruning, leaving the system with fewer inactive synapses. Microglia could strengthen and support active synapses, and lead to a more active TCA cycle in the assembloids. Furthermore, the electrophysiological properties of assembloids suggest a higher excitability of neurons.

### Midbrain-microglia assembloids as a new model for neuro-inflammation in Parkinson’s disease

Microglia play an important role in neurodegenerative diseases. Reactive microgliosis and neuro- inflammation are known to promote neuronal cell death and the pathophysiology of those disorders (Block, Zecca and Hong, 2007; Duffy *et al*., 2018). Midbrain organoids have been used to model neurodegeneration in PD *in vitro* (Kim *et al*., 2019; Smits *et al*., 2019). However, until now, no 3D in vitro model for studying neuro-inflammation in PD was described. The new assembloid model presented here will enable to study reactive microgliosis and neuro- inflammation in PD. Furthermore, a personalized approach, in which patient-specific assembloids are used, can be assessed for genetic and idiopathic cases of PD. This opens doors to studies of neuro-inflammation related pathways and, to new therapeutic targets for compounds that focus on the immune system in the brain.

## Acknowledgments

The authors would like to thank Dr. Jared Sterneckert for providing us with the cells from K7 line, and Dr. Christine Klein from the Institute of Neurogenetics, University of Lübeck for providing the 163 line. J.J. is supported by a Pelican award from the Fondation du Pelican de Mie et Pierre Hippert-Faber. SLN is supported by the National Centre of Excellence in Research on Parkinson’s Disease (NCER-PD) is funded by the Luxembourg National Research Fund (FNR/NCER13/BM/11264123). We thank the LCSB Metabolomics Platform, and specially Christian Jäger and Xiangyi Dong for their contribution to this manuscript. We also thank Dr. Paul Antony and Dr. Silvia Bolognin for the development of the image analysis codes used in this study. We thank Dr. Jennifer Modamio for the development of the R script used for the MEA analysis. The JCS lab is supported by the Fonds National de la Recherche (FNR) Luxembourg (PRIDE17/12244779/PARK-QC; FNR/PoC16/11559169, FNR/ NCER13/BM/11264123). This is an EU Joint Program - Neurodegenerative Disease Research (JPND) project (INTER/JPND/15/11092422). We also would like to thank the private donors who support our work at the Luxembourg Centre for Systems Biomedicine. JWS and JYL are supported by research grant to RIKEN Integrative Medical Sciences (IMS) from the Japanese Ministry of Education, Culture, Sports, Science and Technology (MEXT) and JYL is part of the International Program Associate (IPA) program in RIKEN.

## Resource availability

All original and processed data as well as scripts that support the findings of this study are public available at this https://doi.org/10.17881/cx25-ht49.

## Author contributions

SSS designed and conducted the experiments, interpreted the data and drafted the manuscript. SLN analyzed data. EB, UD, KB, YJL, TK, JJ, GR, Js, HK and CT conducted specific experiments. CS contributed to the manuscript. JWS, SC and JCS coordinated and conceptualized the study. All authors reviewed and approved the final manuscript and agreed to be accountable for their contributions.

## Ethics Statement

Written informed consent was obtained from all individuals who donated samples to this study and all work with human stem cells was done after approval of the national ethics board, Comité National d’Ethique de Recherche (CNER), under the approval numbers 201305/04 and 201901/01.

## Rights retention statement

This research was funded in part by the Fonds National de la Recherche (FNR) Luxembourg. For the purpose of Open Access, the author has applied a CC BY public copyright license to any Author Accepted Manuscript (AAM) version arising from this submission.

