## Supplementary Information for "Microglia integration into human midbrain organoids leads to increased neuronal maturation and functionality"

**Supplemental information**

**Supplemental Figure Legends**

**Figure S1. Microglia in assembloids show different morphologies and allow astrocyte differentiation A.** iPSCs from the line K7 (upper row), 163 (middle row) and EPI (bottom row) stained for the pluripotency markers TRA-1-60, Nanog (left), TRA-1-81, SOX2 (middle), SSEA-4 and Oct-4 (right). **B.** Zymosan and IBA1 staining on microglia from line K7 (top) and 163 (bottom) differentiated for 10 days. **C.** Immunostaining for IBA1, TH and TUJ1 of midbrain organoids and assembloids with microglia from line K7, 163 and EPI upon culture with midbrain organoid (MOm), microglia (MGLm) or co-culture (cc med) media). **D.** MAP2 positive (MAP2^+^) cells in midbrain organoids and assembloids (n (midbrain organoids) =5, 5 batches, n (assembloids) =15, 5 batches, 3 cell lines). For immunofluorescence images see Figure 2C and S1H. Data are represented as mean ± SEM. Y axis is fold change compared to midbrain organoids. **E.** Immunostaining of neural precursor cells from line K7 for the pluripotency marker SOX2, the neural precursor marker Nestin and the forebrain and hindbrain marker PAX6, whose absence confirms the midbrain patterning of the cells. **F.** Bright field images of midbrain organoids (upper panels) and assembloids (bottom panels) at the co-culture day (0DOC, days of co-culture, left) and at 20DOC (right). **G.** Immunostaining of assembloids for IBA1 showing ramified (left), elongated (middle) and round (right) microglia. **H.** Immunostaining of an assembloid from the line 163 after 70 days of co-culture, for GFAP, IBA1 and MAP2. **I.** Immunofluorescence staining of midbrain organoids (left panels) and assembloids with microglia from line 163 (middle panels) and EPI (right panels) for IBA1, FOXA2 and MAP2 (upper panels), and for TH and TUJ1 (bottom panels). The images correspond to a maximum intensity projection from a z-stack of 70µm organoid sections.

**Figure S2. Cell type specific gene expression in assembloids.** Heatmap showing the cell type-specific gene expression throughout the different cell clusters in assembloids.

**Figure S3. Quality control of sn-RNAseq data using the Seurat R Package.** Quality controls of **A.** Midbrain organoids. **B**. Assembloids with microglia from line K7. **C**. Assembloids with microglia from line 163. i) Before quality controls; ii) After quality controls (100 < nFeature_RNA < 5000 or percent_mt < 25); iii) Correlation between features and counts; iv) Correlation between microglial genes and counts. **D.** Volcano plot showing the most variable genes between midbrain organoids and assembloids. **E.** UMAP visualization of scRNA-seq data grouped by samples: Assembloids with microglia from 163 Assembloids_163), Assembloids with microglia from K7 (Assembloids_K7) and midbrain organoids. **F.** Principle components analysis. 20 dimensions were chosen.

**Figure S4. Microglia in assembloids lead to differential expression of multiple genes. A.** Heatmap of the average expression of 100 most significant differentially expressed genes across cell clusters in midbrain organoids and assembloids (p<0.05). **B.** Venn diagrams showing the number of DEG across cell types. Overlap within three neuronal clusters (left pannel) and midNESC, PROG, RGL and NB (right panel). **C.** Complete enrichment analysis of cluster specific DEG between midbrain organoids and assembloids reveals significant network processes (FDR<0.05).

**Figure S5. Microglia lead to a decrease in the inflammasome-related gene PPIA, and differences in synapse-related genes. A.** Expression of PPIA gene, involved in pyroptosis, across cell clusters in midbrain organoids and assembloids. **B.** Expression of *VMAT2* and *SNAP25*, involved in synaptic vesicle exocytosis, across cell clusters in midbrain organoids and assembloids. **C.** Expression levels of axonal guidance and growth-related genes: semaphorins (*SEMA3C*, *DPYSL2*), plexins, ephrins (*EPHA5*), neuropilins, neurofilaments and actin cytoskeleton (*NEFM, ACTB*) across cell clusters in midbrain organoids and assembloids. Data are represented as mean ± SD. * p< 0.05 using a Wilcox test. Dots represent single cells. **D.** Boxplot from patch clamp data showing that neurons in midbrain organoids and assembloids show similar resting membrane potentials (left) and input resistances (middle), and fired repetitive action potentials in response to somatic current injections (right, n midbrain organoids =14; n assembloids = 13 cells). **E.** Inward currents of an assembloid neuron triggered by voltage steps to different potentials starting from -70 mV in voltage-clamp mode. Voltage-gated currents appeared at -40mV and persisted until +30mV under whole-cell voltage-clamp conditions (left). Inward currents generated at -30 mV (red trace in left panel) between midbrain organoids and assembloids showed no significant difference between both groups (right). **F.** Heatmap showing extracellular metabolite levels in culture supernatants (n (midbrain organoids) = 5, 5 batches, n (assembloids) = 15, 5 batches, 3 cell lines). Midbrain organoids cluster separately from assembloids from lines K7, 163 and EPI.

**Figure S6. Assembloids show a different extracellular metabolite profile compared to midbrain organoids. A.** Metabolite levels in the culture media from organoids and assembloids after 48h of culture. The levels of glucose and pyruvic acid were lower in media from assembloids compared to midbrain organoids. **B.** The levels of the amino acids phenylalanine, tyrosine, methionine, lysine, putrescine, threonine, leucine, isoleucine, valine, asparagine and serine in the media were lower in assembloids, whereas the glutamate levels were higher (n (midbrain organoids)= 3, 3 batches, n(assembloids)= 9, 3 batches, 3 cell lines). Each dot represents a replicate (medium 3 organoids or assembloids pooled). Data are represented as mean ± SEM. *p < 0.05, **p<0.01, ***p<0.001, ****p<0.0001 using a Mann-Whitney or an unpaired t test.

**Supplemental video legends**

**Video S1.** Time-lapse of K7 microglia phagocytosing zymosan particles. 449.55 frames, 29.97 frames per second.

**Video S2.** Time-lapse of 163 microglia phagocytosing zymosan particles. 659.34 frames, 29.97 frames per second.

**Video S3.** Time-lapse of EPI microglia phagocytosing zymosan particles. 689.31, 29.97 frames per second.

**Supplemental Tables**

| Name | Simplified identifier | | Identifier | Patient | Gender | Age of sampling | | Source |
| --- | --- | --- | --- | --- | --- | --- | --- | --- |
| K7 | 200 | 2.0.0.10.1.0 | | K7.1 WT/C4 WT | Female | 81 | Reinhardt et al., 2013 | |
| EPI | 201 | 2.0.0.15.0.0 | | A13777 | Female | Cord Blood | GIBCO/A13777 | |
| 163 | 304 | 2.0.0.79.0.0 | | 163 | Male | 66 | SYSMED | |

**Table S1 related to experimental procedures**. iPS cell lines used in this study. Macrophage precursors were derived from iPSCs as described in the Experimental procedures section. Human neural precursors were derived from human iPSCs from the simplified identifier 200. Human midbrain-specific organoids were generated with neural precursors as described in the Experimental procedures section.

| Reagent name | Company | Catalog number | Concentration |
| --- | --- | --- | --- |
| Advanced DMEM/F12 | Thermo Fisher | 12634010 |  |
| N2 | Thermo Fisher | 17502001 | 1x |
| Pen/Strep | Invitrogen | 15140122 | 1x |
| GlutaMax | Thermo Fisher | 35050061 | 1x |
| 2-mercaptoethanol | Thermo Fisher | 31350-010 | 50 μM |
| IL-34 | Peprotech | 200-34 | 100 ng/mL |
| GM-CSF | Peprotech | 300-03 | 10 ng/mL |
| BDNF | Peprotech | 450-02 | 10 ng/mL |
| GDNF | Peprotech | 450-10 | 10 ng/mL |
| DAPT | R&D Systems | 2634/10 | 10 μM |
| Activin A | Thermo Fisher | PHC9564 | 2.5 ng/mL |

**Table S2 related to experimental procedures**. Co-culture medium composition.

| Antibody | Host species | Source | Ref.-No. | Dilution |
| --- | --- | --- | --- | --- |
| IBA1 | Goat | Abcam | ab5076 | 1:250 |
| PU1 | Rabbit | Cell signalling | 2258S | 1:250 |
| CD45 | Mouse | Biolegend | 304002 | 1:1000 |
| TMEM119 | Rabbit | Sigma | HPA051870 | 1:250 |
| P2RY12 | Rabbit | Sigma | HPA014518 | 1:250 |
| FOXA2 | Mouse | Santa Cruz | sc-101060 | 1:250 |
| LMX1A | Rabbit | Abcam | ab139726 | 1:100 |
| PAX6 | Rabbit | Biolegend | 901301 | 1:300 |
| SOX2 | Goat | R&D systems | AF2018 | 1:100 |
| SOX2 | Rabbit | Abcam | ab97959 | 1:100 |
| Nestin | Mouse | Millipore | MAB5326 | 1:100 |
| TH | Rabbit | Abcam | ab112 | 1:1000 |
| TUJ1 | Chicken | Millipore | AB9354 | 1:1000 |
| MAP2 | Chicken | Abcam | ab5392 | 1:1000 |
| MAP2 | Mouse | Millipore | MAB3418 | 1:200 |
| GFAP | Chicken | Millipore | AB5541 | 1:1000 |
| SSEA-4 | Mouse | Millipore | MAB4304 | 1:50 |
| Oct-4 | Rabbit | Abcam | ab19857 | 1:400 |
| TRA-1-60 | Mouse | Millipore | MAB4360 | 1:50 |
| Nanog | Rabbit | Millipore | AB5731 | 1:200 |
| TRA-1-81 | Mouse | Millipore | MAB4381 | 1:50 |
| VAMP2 | Rabbit | Abcam | ab215721 | 1:1000 |
| Actin-β | Mouse | Cell signalling | 3700 | 1:100000 |
| Alexa Fluor® 647 Anti-chicken | Donkey | Jackson Immuno | 703-605-155 | 1:1000 |
| Alexa Fluor® 488 anti-goat | Donkey | Invitrogen | A-11055 | 1:1000 |
| Alexa Fluor® 568 anti-goat | Donkey | Thermo Fisher | a11057 | 1:1000 |
| Alexa Fluor® 647 anti-goat | Donkey | Invitrogen | a21447 | 1:1000 |
| Alexa Fluor® 488 anti-rabbit | Donkey | Thermo Fisher | a21206 | 1:1000 |
| Alexa Fluor® 568 anti-rabbit | Donkey | Invitrogen | a10042 | 1:1000 |
| Alexa Fluor® 647 anti-rabbit | Donkey | Invitrogen | a31573 | 1:1000 |
| Alexa Fluor® 488 anti-mouse | Donkey | Invitrogen | a21202 | 1:1000 |
| Alexa Fluor® 568 anti-mouse | Donkey | Invitrogen | a10037 | 1:1000 |
| Hoechst 33342 Solution (20 mM) |  | Invitrogen | 62249 | 1:10000 |
| IgG H+L 800 | Rabbit | Cell signalling | 5151 | 1:10000 |
| IgG H+L 680 | Mouse | Cell signalling | 5470 | 1:10000 |

**Table S3 related to experimental procedures**. Antibodies used in this study.

| Primer | Sequence (5’ to 3’) | Region (Purpose) |
| --- | --- | --- |
| h-RPL37A-F | GTGGTTCCTGCATGAAGACAGTG | RT-PCR |
| h-RPL37A-R | TTCTGATGGCGGACTTTACCG | RT-PCR |
| 2241_AIF1_F | AGACGTTCAGCTACCCTGACTT | RT-PCR |
| 2242_AIF1_R | GGCCTGTTGGCTTTTCCTTTTCTC | RT-PCR |
| 2243_CD68_F | CTTCTCTCATTCCCCTATGGACA | RT-PCR |
| 2244_CD68_R | GAAGGACACATTGTACTCCACC | RT-PCR |
| 2289_P2RY12 FW | AAGAGCACTCAAGACTTTAC | RT-PCR |
| 2290_P2RY12 RV | GGGTTTGAATGTATCCAGTAAG | RT-PCR |
| 2291_TMEM119 FW | AGTCCTGTACGCCAAGGAAC | RT-PCR |
| 2292_TMEM119 RV | GCAGCAACAGAAGGATGAGG | RT-PCR |

**Table S4 related to experimental procedures**. Primers used in this study.

| **Stem**  **cells** | **DA** | **VTA** | **Maturity** | **CN** | **GAN** | **GLN** | **SN** |
| --- | --- | --- | --- | --- | --- | --- | --- |
| SOX2  PAX6  HES5  ASCL1  SOX1  PAX3  DACH1  LMO3  NR2F1  PLAGL1  LIX1  HOXA2  FOXA2  SLC1A3  MSI1  VIM  NES  SHH | NR4A2  PBX1  GRIA3  TH  EN1  TMCC3  NTM  DDC  CAMK2N1  ALDH1A1  APP  PDZRN4  PCDH10  ERBB4  SLC10A4  BEX5  NPY1R  GPC2  HTRA2  HTRA3  HTRA4  PTX3  OTX2  EN2  SOX6  FOXA2  LMX1A  LMX1B  KCNJ6  CALB1  SLC6A3  SHH  NKX6−1 | ALDH1A1  TRHR  CD24  SLC18A2  FGF1  NRIP3  MPP6  NTS  CCK  SOX6  GRIN2C  SNCG  IGF1  ADCYAP1  GRP  LPL  CALB1  SLC32A1  VIP  TACR3  DCC  OTX2  SATB1 | SPRYD7  TUBB2A  SKP1  GNAI1  MRAS  ATP1A3  MAGED1  ARL2  MCFD2  MORF4L2  NRSN2  NAP1L3  NGRN  OLFM1  DKK3  CCDC136  COX6C  HSP90AB1  CALM2  ATP1B1  UQCRB  COX4I1  PSMB5  FXYD7  RTN1  SEC62  COX7C  CNTN1  FAIM2  SLC48A1  RAB3B  NPTXR  PDGFA  NDUFC2  BEX2  UCHL1 | CHAT  SLC18A3  ACHE | GAD1  GAD2  GABARAP  GABARAPL1  GABARAPL2  ABAT | SLC1A1  SLC1A2  SLC1A3  SLC17A6  SLC17A7  GLS  GLS2  GRIN1  GRIN2A  GRIN2B  GRIN2C  GRIN2D  GRIN3A  GRIN3B  GRINA  GRIA1  GRIA2  GRIA3  GRIA4  HTRA1 | SLC6A4  SLC18A2  TPH1  TPH2  FEV  HTR1D  HTR1E  HTR1F  HTR2A  HTR2A−AS1  HTR2B  HTR2C  HTR3A  HTR3B  HTR3D  HTR4  HTR5A  HTR5A−AS1 |

**Table S5 related to results.** Genes used for defining neuronal cell clusters and maturity. DA = Dopaminergic; VTA = ventral tegmental area; CN = Cholinergic; GAN = Gabaergic; GLN = Glutamatergic; SN = Serotonergic.

| Name | Identifier |
| --- | --- |
| Cell 1 | MglK7_CATCAGAAGGCCATAG.1 |
| Cell 2 | MglK7_AGGGATGGTGTCCTCT.1 |
| Cell 3 | MglK7_CTAAGACGTCTAGCGC.1 |
| Cell 4 | MglK7_CCGTGGACATCCTTGC.1 |
| Cell 5 | MglK7_AGTTGGTCAGCCTTTC.1 |
| Cell 6 | Mgl163_ACGCCAGTCATTGCCC.1 |
| Cell 7 | Mgl163_CATATTCTCCAATGGT.1 |
| Cell 8 | MglK7_CGCTGGAGTCCAGTTA.1 |
| Cell 9 | MglK7_AAGCCGCAGCTTCGCG.1 |
| Cell 10 | MglK7_CTCCTAGTCCTGCTTG.1 |
| Cell 11 | MglK7_CAAGGCCGTAGGCATG.1 |
| Cell 12 | MglK7_ACAGCCGCAGCTGCAC.1 |
| Cell 13 | MglK7_CAACCAACAGGAATGC.1 |
| Cell 14 | MglK7_ACGGGCTGTACAAGTA.1 |
| Cell 15 | MglK7_GTCTTCGCAATCCGAT.1 |
| Cell 16 | Mgl163_GCACATACATCGGACC.1 |
| Cell 17 | Mgl163_AGATCTGCAAACGTGG.1 |
| Cell 18 | Mgl163_ACGTCAAAGTTTAGGA.1 |
| Cell 19 | MglK7_GGATGTTGTCGCCATG.1 |
| Cell 20 | MglK7_TGTTCCGCAAGAGGCT.1 |
| Cell 21 | MglK7_GACTACAAGTAGGCCA.1 |
| Cell 22 | MglK7_TTAGGACAGGTTACCT.1 |
| Cell 23 | MglK7_GCTGGGTAGCGATATA.1 |
| Cell 24 | MglK7_GCTTGAAAGATACACA.1 |
| Cell 25 | MglK7_AAAGTAGGTGACTACT.1 |
| Cell 26 | MglK7_TATCTCACAGGTTTCA.1 |

**Table S6 related to results**. Identifiers of the 26 microglia cells.
